# SGLT2 Inhibitors Rescue Lysosomal mTORC1 Hyperactivity and Proximal Tubulopathy in Preclinical Models of Cystinosis

**DOI:** 10.64898/2026.01.29.702541

**Authors:** Svenja Aline Keller, Patrick Krohn, Zhiyong Chen, Alessandro Luciani, Olivier Devuyst

## Abstract

The loss of lysosomal cystine transporter cystinosin (CTNS) disrupts kidney proximal tubule (PT) function, causing cystinosis - a prototypical lysosomal storage disorder characterized by cystine accumulation and metabolic dyshomeostasis. Cystine storage disrupts lysosomal nutrient sensing and downstream mTORC1 signalling, driving loss of PT differentiation and proximal tubulopathy. Here, using cross-species disease models, differentiated cellular systems, lysosome-based assays, and transcriptomics profiling, we demonstrate that sodium-glucose co-transporter 2 (SGLT2) inhibitors (empagliflozin or dapagliflozin) ameliorate proximal tubulopathy in cystinosis. In CTNS-deficient PT cells, SGLT2 inhibition restores lysosome proteolysis, autophagic flux, metabolic homeostasis, and epithelial differentiation and function, independently of cystine clearance. Mechanistically, SGLT2 inhibition reduces the assembly of the v-ATPase-Ragulator-Rag scaffolding complex at lysosomes, thereby decoupling cystine storage from pathological mTORC1 activation. These effects reprogram PT metabolic trajectories and differentiation states, mitigating proximal tubulopathy across zebrafish and rodent models of cystinosis. Together, these findings define a lysosome-metabolism crosstalk that links glucose handling to mTORC1 regulation and provide a rationale for repurposing SGLT2 inhibitors as a disease-modifying therapy for cystinosis and related lysosome-driven PT disorders.

## INTRODUCTION

Cystinosis is a prototypical lysosomal storage disease (LSD) caused by inactivating mutations in the *CTNS* gene, which encodes cystinosin, a proton-driven cystine exporter at the lysosomal membrane.^1,2^ Loss of CTNS leads to intralysosomal cystine accumulation across tissues and a multisystem disorder with early and prominent kidney involvement. Nephropathic cystinosis (OMIM #219800), the most severe and common form of cystinosis, manifests in infancy as early-onset proximal tubulopathy that progresses to chronic kidney disease (CKD) and systemic metabolic complications.^3,4^ Cysteamine — the only approved therapy — reduces lysosomal cystine burden and delays progression, but it does not prevent proximal tubular dysfunction and is limited by tolerability and adherence.^5^ Gene therapy replacement strategies have shown promise in preclinical and early clinical settings, but remain constrained by cost, technical complexity, and accessibility.^6,7^ Together, these limitations underscore the need for safe, affordable, and widely deployable therapies that preserve PT function downstream of cystine storage.

The PT is a highly specialized epithelial segment that relies on receptor-mediated endocytosis and lysosomal processing to reclaim ultrafiltered low-molecular-weight (LMW) proteins and solutes. Recent work, including ours, has shown that cystine storage in CTNS-deficient cells disrupts lysosome-based nutrient sensing, leading to aberrant activation of mechanistic target of rapamycin complex 1 (mTORC1) via the Ragulator–Rag GTPase machinery.^8^ Sustained lysosome-associated mTORC1 activation reprograms PT cells toward anabolic states at the expense of their differentiated, reabsorptive phenotype, impairing endocytosis and causing low-molecular-weight (LMW) ultrafiltered proteins, resulting in LMW proteinuria – the earliest and most consistent clinical hallmark of nephropathic cystinosis.^8–10^

Pharmacologic inhibition of mTORC1 can rescue autophagy-lysosome defects and improve PT differentiation and function in preclinical cystinosis models, supporting lysosome-associated mTORC1 as a key downstream pathogenic node.^8–10^ However, clinical translation of direct mTOR inhibitors is limited by systemic toxicity and incomplete pathway selectivity, motivating alternative strategies that normalize — rather than ablate — pathological mTORC1 activation downstream of lysosomal dysfunction.^11,12^

Targeting metabolic inputs upstream of mTORC1 is an attractive approach in that context.^13^ In the kidney, the sodium-glucose co-transporter 2 (SGLT2 encoded by *SCL5A2*) is highly expressed at the apical membrane of PT cells and mediates the bulk of filtered glucose reabsorption.^14^ SGLT2 inhibitors (SGLT2i), developed for diabetes, provide robust kidney-protective effects across diabetic and non-diabetic kidney disorders, with a favourable safety profile.^15,16^ Beyond glucosuria, SGLT2i induce coordinated metabolic adaptations in PT cells that alter substrate availability and energy balance and can secondarily modulate lysosome-centered signaling pathways, including mTORC1 and autophagy–lysosome function.^17,18^

Human genetics supports the tolerability of sustained SGLT2 attenuation. Individuals with SLC5A2 loss-of-function (familial renal glucosuria) have lifelong glycosuria with preserved kidney function, indicating that chronically reduced PT glucose flux is generally compatible with metabolic adaptation rather than injury.^19^ Stress-associated ketosis (e.g., fasting or pregnancy) further suggests metabolic flexibility. Together with evidence that limiting lysosome-centered nutrient signalling suppresses pathological mTORC1 activity and promotes epithelial differentiation independently of ketogenesis,^8,20^ these findings support the idea that SGLT2 inhibition can restore PT homeostasis downstream of cystine storage.

Here, we integrate cross-species cystinosis models, physiologically relevant PT systems, lysosome-centered functional assays, and transcriptomics to define the consequences of SGLT2 inhibition. We show that SGLT2i restore lysosome-associated mTORC1 activity toward wild-type levels, re-engage autophagy and metabolic homeostasis, and rescue PT differentiation and endocytic function, mitigating proximal tubulopathy downstream of CTNS loss and cystine storage. These findings link PT glucose flux to lysosome-centered mTORC1 regulation and provide a mechanistic rationale for repurposing SGLT2i as disease-modifying therapies for cystinosis and related mTORC1-driven proximal tubulopathies.

## RESULTS

### SGLT2 inhibition mitigates hyperactive mTORC1 and rescues a zebrafish model of cystinosis

We evaluated the therapeutic potential of SGLT2i in a zebrafish model of cystinosis, which provides a sensitive and physiologically relevant platform for phenotypic drug screening.^21,22^ Consistent with disease progression, *ctns* mutant larvae accumulated cystine, exhibited hyperactive mTORC1 activity detectable by 5 days post-fertilization (dpf), and developed defective endocytosis and lysosome-related pathways with LMW proteinuria by 14 dpf (**Figure S1A-S1E**).

The *ctns*-deficient larvae expressed *slc5a2* in the pronephros at levels comparable to wild-type larvae (**<u>Figure S1F and S1G</u>**). Treatment with low, non-toxic doses of dapagliflozin or empagliflozin (**<u>Figure S1H and S1I</u>**, ref. ^23^) reduced mTORC1 activity in *ctns* mutants toward wild-type levels (**Figure 1A and 1B**). By comparison, rapamycin suppressed mTORC1 activity below wild-type levels (**Figure 1B**).^13^ Using *lfabp::½vdbp–mCherry* zebrafish as a *bona fide* reporter of PT endocytic function,^22^ SGLT2 inhibition improved lysosomal processing of ultrafiltered LMW proteins, reflected by reduced pronephric mCherry signal (**Figure 1C**). Consistently, urinary LMW protein excretion decreased dose-dependently in SGLT2i-treated *ctns* mutants compared with vehicle- or cysteamine-treated mutants (**Figure 1D**). Notably, these improvements occurred without detectable changes in whole-larva cystine levels (**Figure 1E**), indicating that SGLT2 inhibition ameliorates proximal tubulopathy downstream of ctns deficiency and cystine storage.

**Figure 1.**
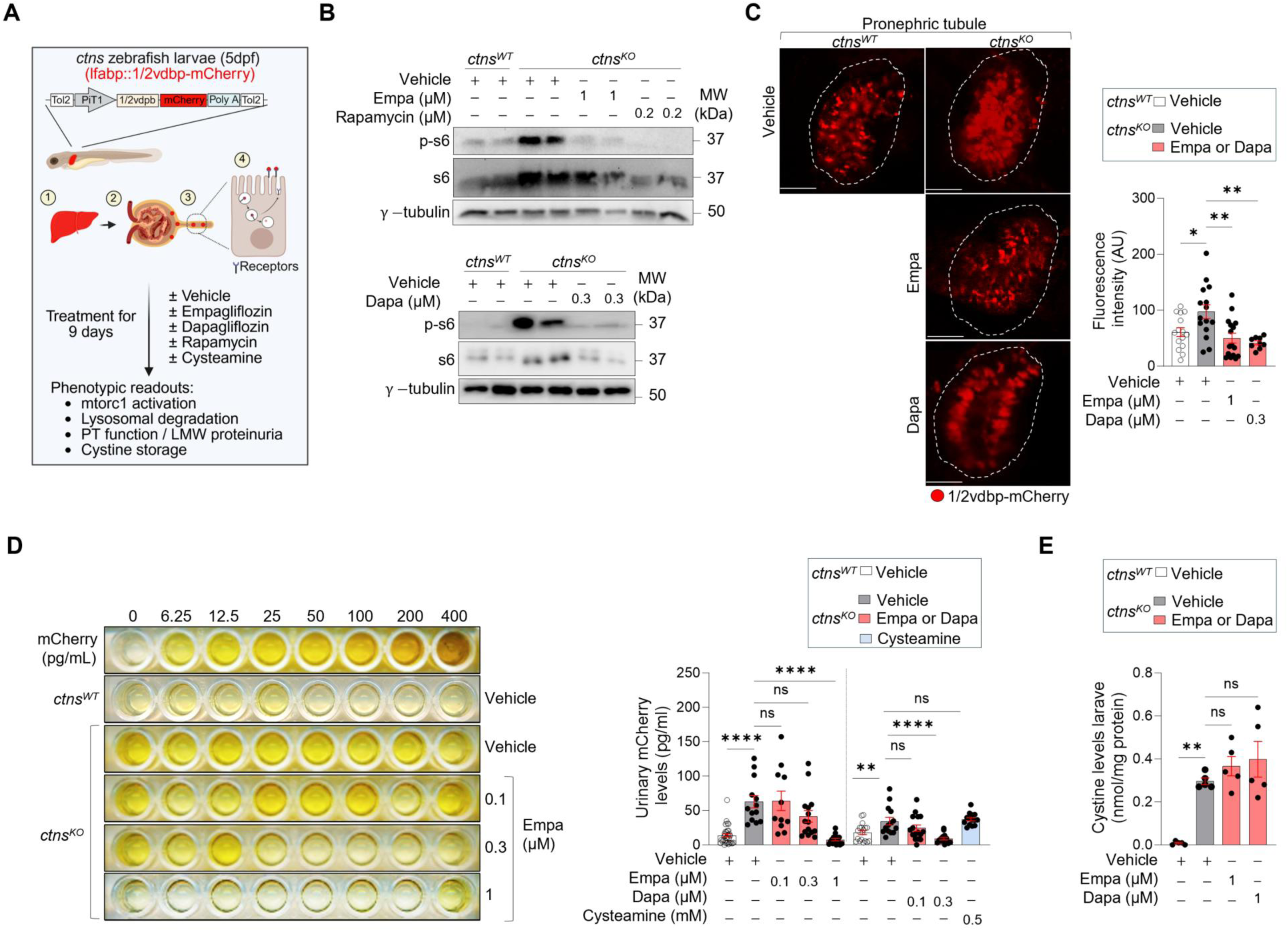
SGLT2 inhibition reduces mTORC1 activity toward wild-type levels and mitigates LMW proteinuria in a zebrafish model of cystinosis. (**A**) Schematic of the *ctns* zebrafish line expressing the vdbp–mCherry reporter and treatment regimen. vdbp–mCherry (∼50 kDa) is synthesized in the liver, released into the circulation, filtered by the glomerulus, taken up by PT cells via receptor-mediated endocytosis, and degraded in the endolysosomal system. *ctns* larvae expressing vdbp–mCherry were treated from 5dpf with vehicle, empagliflozin (empa), dapagliflozin (dapa), rapamycin, or cysteamine for 9 days. (**B**) Immunoblotting of p-s6/s6. Each lane represents a pool of 6 zebrafish larvae (n = 2 biologically independent pools per condition). (**C**) Multiphoton microscopy and quantification of mCherry fluorescence in the pronephros (n ≥ 15 zebrafish larvae per condition). (**D**) Urinary mCherry levels by ELISA (representative plate image and quantification; n ≥ 12 zebrafish larvae per condition). (**E**) Cystine levels. Each dot represents a pool of 6 zebrafish larvae (n = 5 biologically independent pools per condition). Data are mean ± SEM. Statistical analyses were calculated by one-way ANOVA followed by Sidak’s multiple comparisons test in (B-E). *p < 0.05, **p < 0.01, ****p < 0.0001 versus vehicle-treated wild-type or vehicle-treated *ctns*^KO^ zebrafish. NS, non-significant. Scale bars, 20µm.

### Mechanisms of mTORC1 and homeostasis regulation by SGLT2 inhibition

To define how SGLT2 inhibition attenuates pathological mTORC1 activation downstream of CTNS deficiency, we analyzed primary mouse proximal tubule cells (mPTCs) microdissected from *Ctns^KO^* kidneys. These cells recapitulate key cystinosis hallmarks, including cystine storage, defective lysosomal degradation, impaired autophagy flux, anabolic growth, and loss of PT differentiation and epithelial markers.^8–10^ Given its established pediatric safety, we focused on dapagliflozin for mechanistic studies (**Figure 2A**).^24^

**Figure 2.**
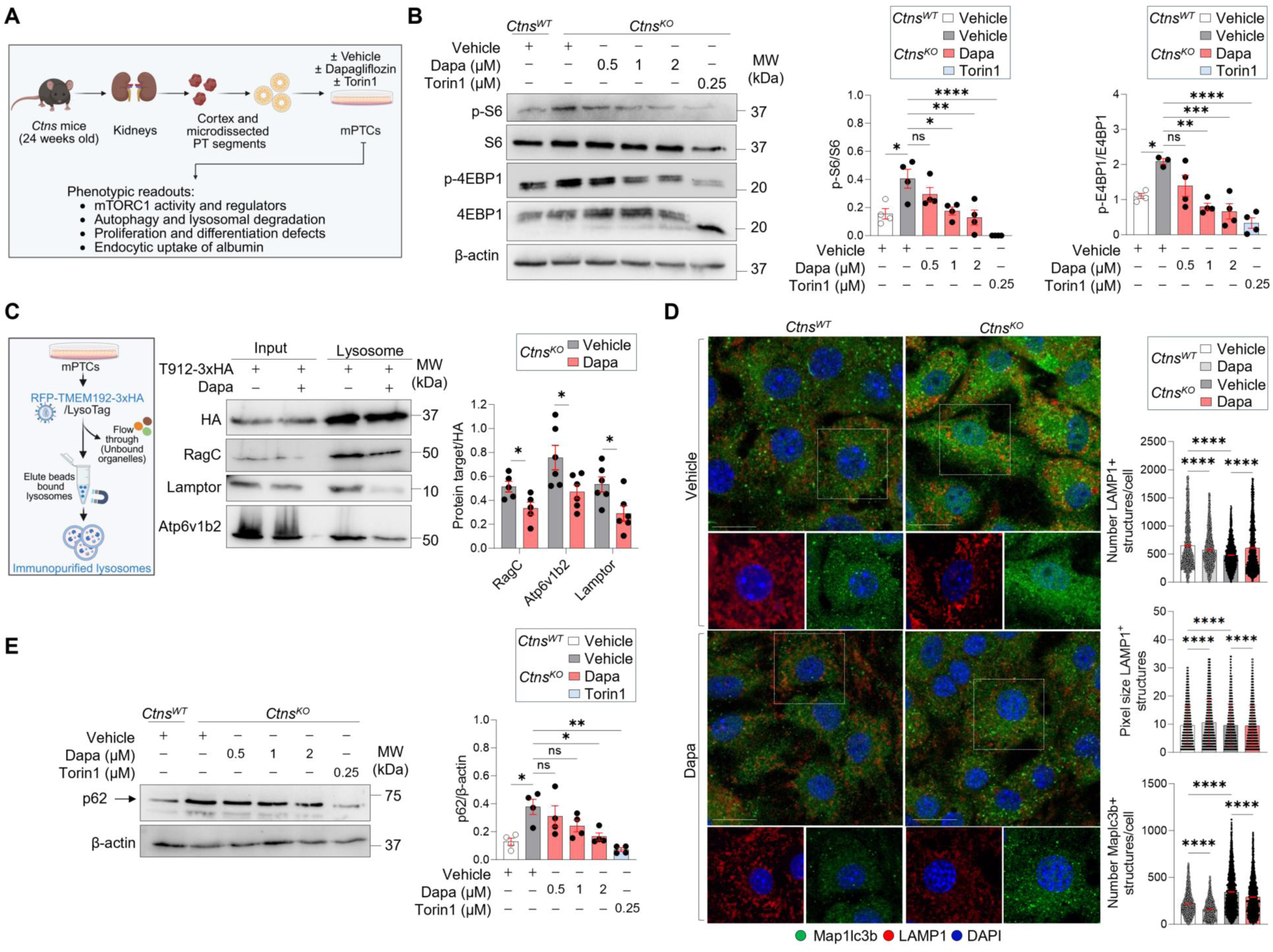
Dapagliflozin reduces mTORC1 signalling and improves lysosome/autophagy readouts in CTNS-deficient PT cells. (**A**) Workflow for the isolation of primary PT cells (mPTCs) from *Ctns* mice. Cells were treated with vehicle (DMSO), dapagliflozin (0.5-2µM) or Torin1 (250nM) for 16h. (**B**) Immunoblotting and quantification of p-S6/S6 and p-4EBP1/4EBP1 (n = 4 biologically independent samples). (**C**) Immunoblotting and quantification of the indicated proteins in input samples and corresponding lysosome-immunopurified (Lyso-IP; 3×HA-tagged TMEM192) fractions from *Ctns*^KO^ mPTCs (n = 5 biologically independent samples). (**D**) Representative confocal micrographs and quantification of Map1lc3b^+^ structures (green) and LAMP1^+^-positive lysosomes (red) (n = 3 biologically independent samples). (**E**) Immunoblotting and quantification of p62 (n = 4 biologically independent samples). Data are mean ± SEM. Statistical analyses were performed using one-way ANOVA followed by Sidak’s multiple comparisons test in (**B, D** and **E**) and unpaired two-tailed Student’s t test in (**C**). *p < 0.05, **p < 0.01, ***p < 0.001, ****p < 0.0001 versus vehicle-treated wild-type or vehicle-treated *Ctns*^KO^ cells. Nuclei counterstained with DAPI (blue). Dotted white squares contain images at high magnification. NS, non-significant. Scale bars, 20µm.

Treatment with non-toxic doses of dapagliflozin (**<u>Figure S2A</u>**) normalized mTORC1 signalling in *Ctns*^KO^ mPTCs in a dose-dependent manner, decreasing the phosphorylation of ribosomal S6 (pS6^Ser2^^35/236^) and 4E-BP1 (p4EBP1^Ser65^) toward wild-type levels (**Figure 2B**). In contrast, the mTOR kinase inhibitor Torin1 resulted in profound suppression of mTORC1 activity below physiological levels (**Figure 2B**). Dapagliflozin reduced mTORC1 signalling despite lower SGLT2 expression in *Ctns*^KO^ mPTCs (**<u>Figure S2B</u>**), indicating that reduced transporter expression does not preclude a measurable pharmacologic response in this system.

At the lysosomal level, dapagliflozin reduced lysosomal enrichment of nutrient-sensing components, including Atp6v1b2, Lamtor2, RagC, consistent with decreased assembly of the v-ATPase-Ragulator-Rag platform (**Figure 2C<u>; Figure S2C</u>**). This was accompanied by improved lysosomal integrity and proteolytic readouts, including increased numbers of morphologically intact LAMP1^+^ lysosomes and enhanced Pepstatin A signal (**Figure 2D<u>; Figure S2D</u>**). Dapagliflozin also improved autophagy-associated markers, with reduced p62/SQSTM1 accumulation and normalization of Map1lcb3^+^ puncta relative to vehicle-treated mutant cells (**Figure 2D and 2E**).

Consistent with recovery of lysosome-autophagy pathways, dapagliflozin shifted PT cells toward a more differentiated profile (**Figure 3A-3B**). Transcript analysis showed increased *Lrp2* and *Cubn* expression and decreased expression of cell-cycle–associated genes (e.g., *Cdc20*, *Ccna2*, *Ccnb2*) (**Figure 3B**). BrdU incorporation confirmed fewer proliferating cells following dapagliflozin treatment (**Figure 3C**). Additional differentiation-associated features improved, including increased primary cilium length (acetylated α-tubulin staining; **Figure 3D**). Functional recovery of endocytic capacity was supported by restored uptake of fluorescently labelled albumin (BSA-A488) (**Figure 3E**).

**Figure 3.**
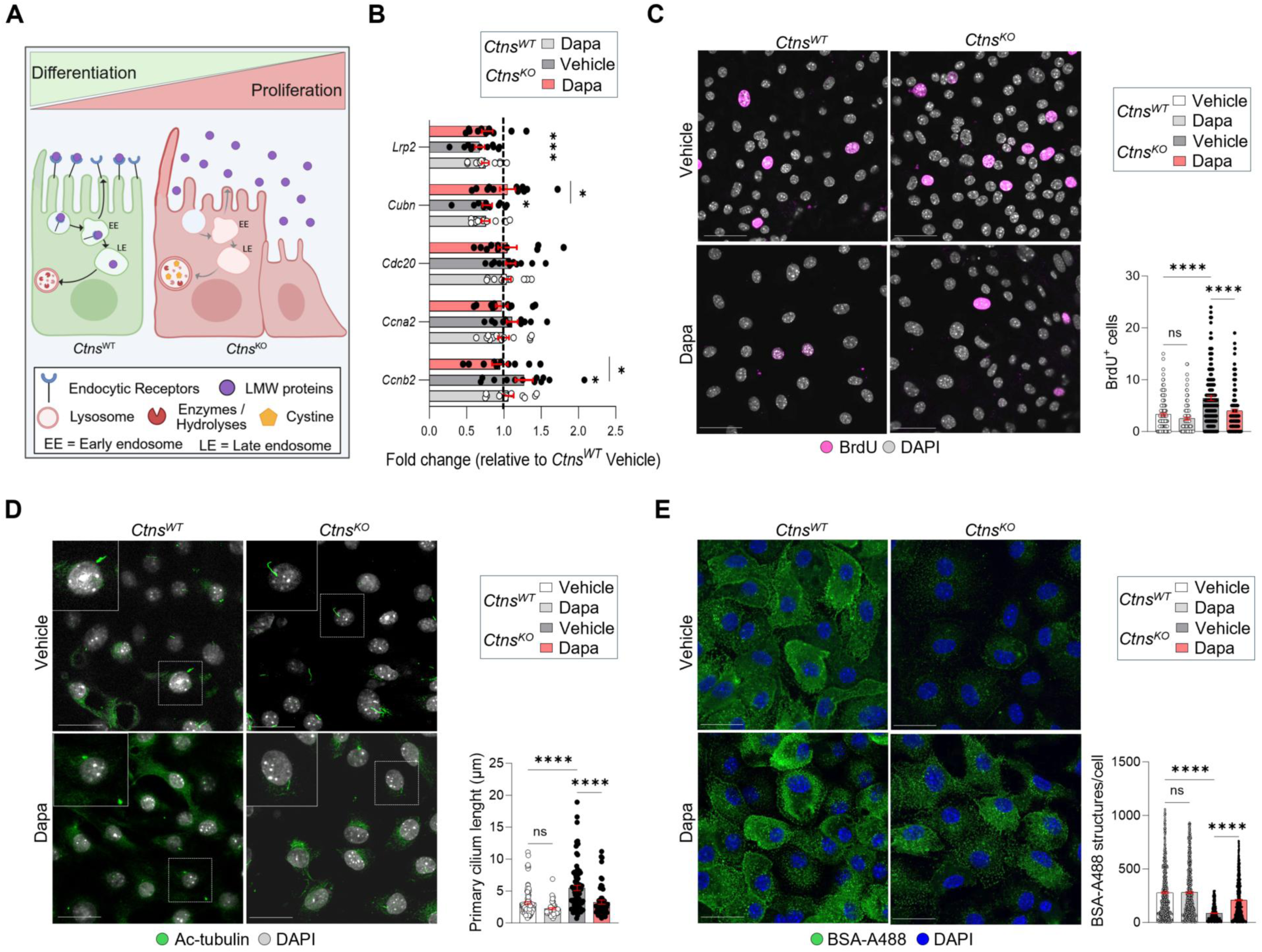
Dapagliflozin improves differentiation and endocytic function in CTNS-deficient proximal tubule cells. (**A**) Schematic summarizing differentiation and proliferation defects in CTNS-deficient PT cells. (**B**) mRNA expression levels of the indicated genes. Gene expression was normalized to *Gapdh* and expressed relative to *Ctns^WT^* vehicle (dotted line, n = 3 biologically independent samples, with n = 4 technical replicates per mouse and condition). (**C**) Proliferation analysis following bromodeoxyuridine (BrdU) incorporation (1.5µg mL⁻¹ for 16 h at 37 °C). Cells were analyzed by confocal microscopy and quantified as the number of BrdU-positive nuclei per field (n = 10 fields pooled from n = 3 biologically independent samples). (**D**) Representative confocal micrographs and quantification of primary cilium length (acetylated tubulin, green; n = 3 fields pooled from n = 3 biologically independent samples). (**E**) Pulse-chase uptake of Alexa 488-BSA (0.2mg/ml): 1h binding at 4°C followed by internalization at 37°C. Representative confocal micrographs and quantification of BSA+ (green) (n = 10 fields pooled from n = 3 biologically independent samples). Data are presented as mean ± SEM. Statistical analyses were performed using an unpaired two-tailed Student’s t test in (**B**) or by one-way ANOVA followed by Sidak’s multiple comparisons test in (**C**-**E**). *p < 0.05, ***p < 0.001, ****p < 0.0001 versus vehicle-treated wild-type or vehicle-treated *Ctns*^KO^ cells. Dotted white boxes indicate regions shown at higher magnification. NS, non-significant. Nuclei counterstained with DAPI (blue or grey, where indicated). Scale bars, 50µm in (**c**), 20µm in (**D**-**E**).

Together, these data show that SGLT2 inhibition reduces pathological mTORC1 signalling and is associated with improvements in lysosome/autophagy readouts and PT differentiation and function in CTNS-deficient mPTCs.

### Therapeutic validation of Dapagliflozin in a rat model of cystinosis

To test SGLT2 inhibition *in vivo*, we evaluated dapagliflozin in *Ctns*^KO^ rats, which recapitulate lysosomal storage and early-onset PT dysfunction characteristic of cystinosis.^10^ Dapagliflozin was administered in drinking water (10 mg/kg/day) for 4 weeks starting at 10 weeks of age, a stage preceding overt proximal tubulopathy but coinciding with early lysosome-associated mTORC1 hyperactivation (**Figure 4A**; ref.^8,10^).

**Figure 4.**
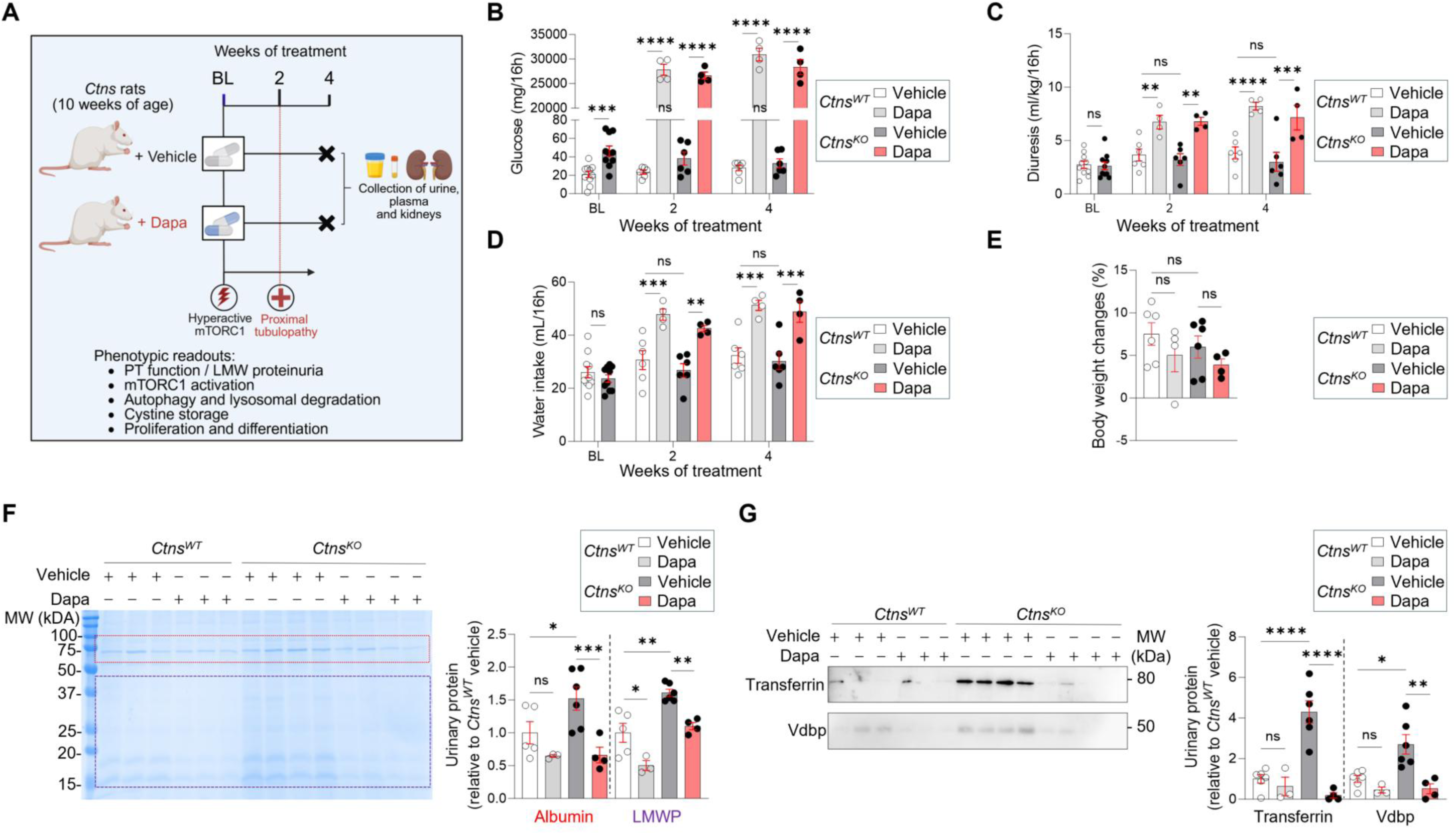
Dapagliflozin mitigates early proximal tubule dysfunction in a rat model of cystinosis. (**A**) Study design. Ten-week-old *Ctns*^WT^ and *Ctns*^KO^ rats received dapagliflozin (10 mg kg⁻¹ day⁻¹) in drinking water for 4 weeks. Urine was collected every 2 weeks; plasma and kidneys were collected at study end. (**B**) Urinary glucose excretion over treatment. (**C**) Overnight urine output (mL per 16h), normalized to body weight. (**D**) Water intake (mL per 16h). (**E**) Body weight changes relative to baseline. (**F**) Coomassie-stained SDS-PAGE analysis of urinary proteins at endpoint and quantification of albumin and LMWP (MW ≤ 50 kDa) (n ≥ 4 rats per genotype and condition). (**G**) Urinary transferrin and Vdbp by immunoblot with densitometry at endpoint. n ≥ 4 rats per genotype and condition. Data are mean ± SEM. Statistical analyses were performed using one-way ANOVA followed by Sidak’s multiple comparisons test in (**B**-**G**). *p < 0.05, **p < 0.01,***p < 0.001, ****p < 0.0001 versus vehicle-treated wild-type or vehicle-treated *Ctns*^KO^ rats. NS, non-significant.

As expected, dapagliflozin induced glucosuria and diuresis, reflected by increased urine volume and water intake (**Figure 4B–4D<u>; Table S1</u>**), despite mildly reduced SGLT2 expression in mutant PTs (**<u>Figure S3A and S3B</u>**). Urinary sodium excretion increased, and plasma glucose modestly decreased (**<u>Tables S1 and S2</u>**). Treated rats also exhibited a mild reduction in body weight (**Figure 4E**).

Functionally, dapagliflozin reduced urinary albumin and LMW protein excretion (**Figure 4F**). Immunoblotting confirmed reduced urinary vitamin D-binding protein (Vdbp) and transferrin, consistent with improved receptor-mediated PT protein reabsorption (**Figure 4G**). These functional changes occurred alongside a modest increase in blood urea nitrogen (BUN) (**<u>Table S2</u>**), which may reflect diuresis and associated volume effects.^25^ Together, these findings indicate that dapagliflozin improves early proximal tubular dysfunction in *Ctns*^KO^ rats.

### Dapagliflozin couples reduced mTORC1 signalling with metabolic and differentiation programs in vivo

We next assessed whether improved PT function in dapagliflozin-treated rats was associated with changes in mTORC1 activity and metabolic state. Immunoblotting and immunofluorescence analyses showed reduced mTORC1 signalling (decreased pS6) and changes consistent with improved autophagy-lysosome function, including increased cathepsin D (CtsD) maturation and reduced p62 accumulation in Aqp1^+^ PT cells (**Figure 5A** **and 5B**).

**Figure 5.**
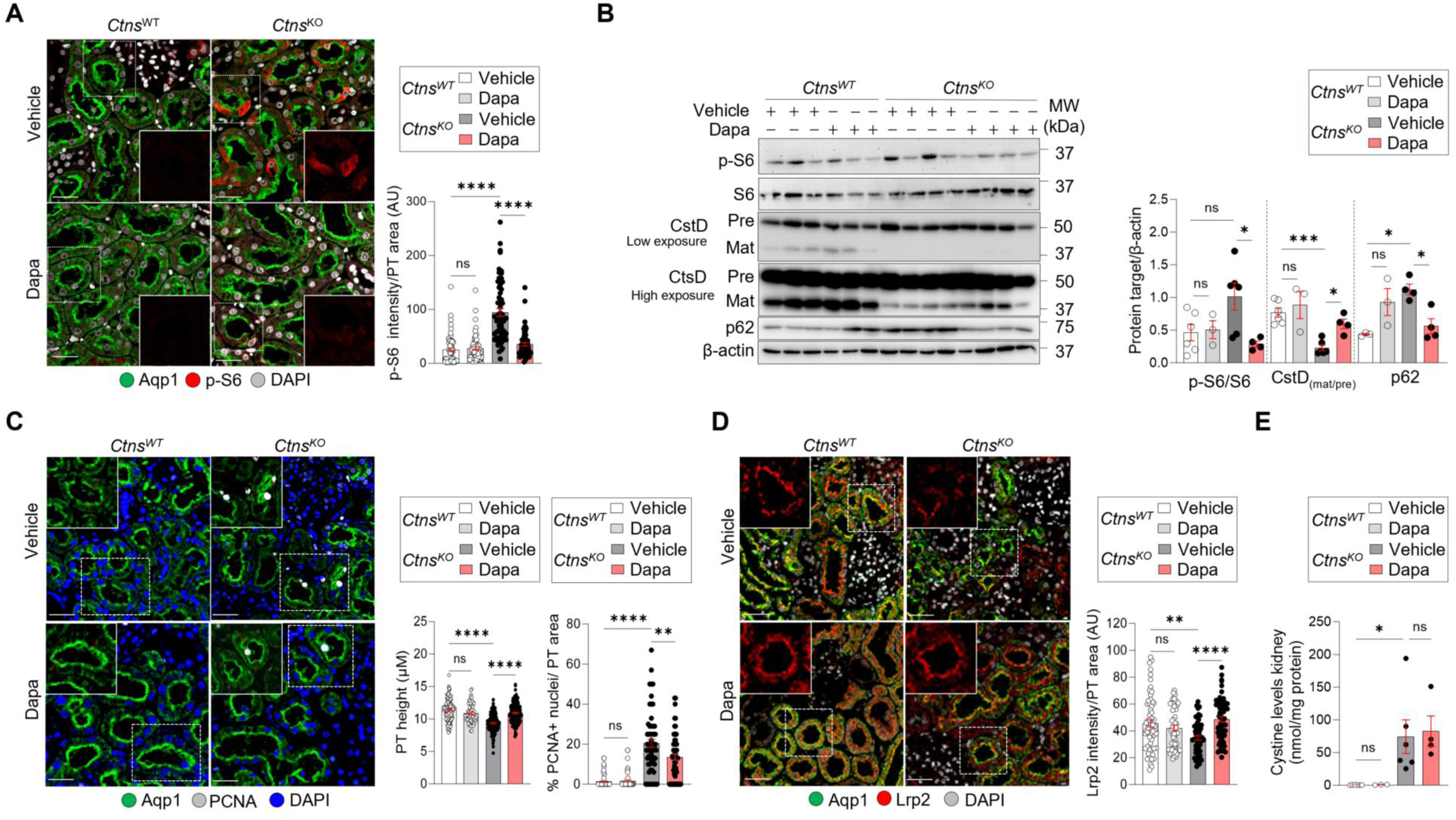
Dapagliflozin is associated with improved lysosome/autophagy readouts and PT differentiation in CTNS-deficient rats. (**A**) Representative confocal micrographs and quantification of p-S6 (red) in the PT after 4 weeks of treatment (n = 3 fields pooled from n = 3 biologically independent samples). (**B**) Immunoblotting and quantification of p-S6/S6, p62 and Cathepsin D (CtsD) after 4 weeks of treatment (n ≥ 4 rats per genotype and condition). (**C**) Representative confocal micrographs and quantification of the height of AQP1^+^ tubules (green) and the number of PCNA^+^ nuclei (white) after 4 weeks of treatment (n = 3 fields pooled from n = 3 biologically independent samples per condition). (**D**) Representative confocal micrographs and quantification of Lrp2 (red) (n = 3 fields pooled from n = 3 biologically independent samples). (**E**) Cystine levels in the kidneys of wild-type and *Ctns*^KO^ rats after 4 weeks of treatment (n ≥ 4 rats per genotype and condition). Data are mean ± SEM. Statistical analyses were performed using one-way ANOVA followed by Sidak’s multiple comparisons test in (**A**-**E**). *p < 0.05, **p < 0.01,***p < 0.001, ****p < 0.0001 versus vehicle-treated wild-type or vehicle-treated *Ctns*^KO^ rats. NS, non-significant. Dotted white boxes indicate images at higher magnification. Nuclei counterstained with DAPI (blue or grey, where indicated). Scale bars, 50µm.

Dapagliflozin treatment was also associated with reduced apoptosis (cleaved caspase-3), decreased vimentin expression, and reduced fibrotic remodelling markers (**<u>Figure S4A–S4D</u>**). These changes were accompanied by improved PT morphology (increased epithelial height), reduced aberrant proliferation (PCNA; **Figure 5C**), and restored apical Lrp2 expression (**Figure 5D**). These effects occurred without changes in kidney cystine levels (**Figure 5E**).

To define transcriptional programs associated with treatment, we performed bulk RNA-seq on whole-kidney lysates from vehicle-treated *Ctns*^WT^ and *Ctns*^KO^ rats and dapagliflozin-treated *Ctns*^KO^ rats. Principal component analysis showed separation by genotype and treatment (**Figure S5A and S5B**). Dapagliflozin shifted the expression of a subset of genes toward the wild-type profile (**<u>Figure S5C</u>**).

Vehicle-treated *Ctns*^KO^ versus *Ctns*^WT^ kidneys showed 1,739 differentially expressed genes (DEGs; 1,020 upregulated, 719 downregulated; |log2FC| > 0.5, P < 0.05; **<u>Figure</u> <u>S5D</u>**). Enrichment analyses identified immune/inflammatory activation, altered transport programs, and broad metabolic dysregulation (**<u>Figure S5E</u>**, ref. ^8–10^). In *Ctns*^KO^ kidneys, dapagliflozin altered 527 genes (**Figure 6A**) and enriched oxidative metabolic pathways, including fatty acid metabolism, the TCA cycle, and gluconeogenesis (**Figure 6B**).

**Figure 6.**
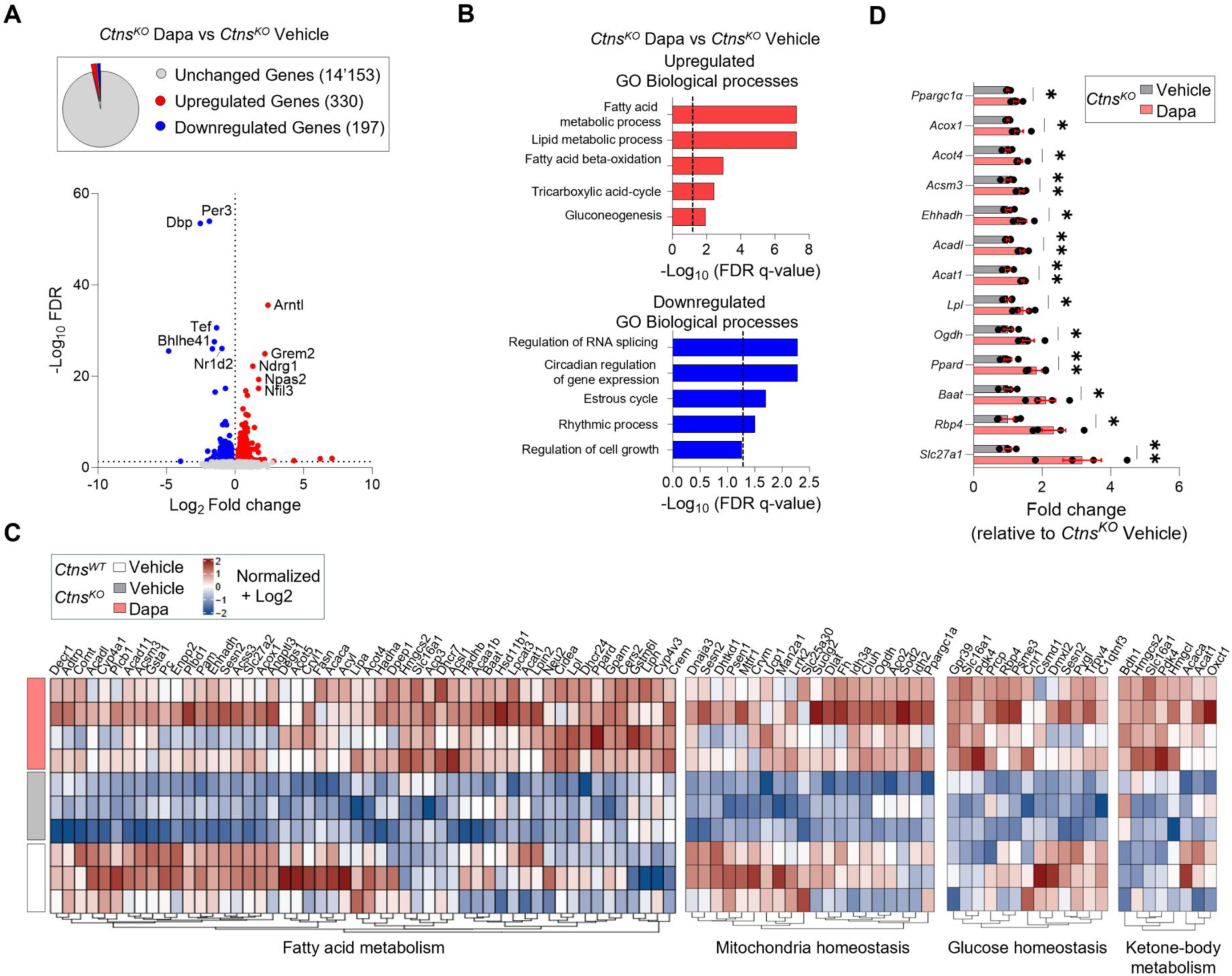
Dapagliflozin induces metabolic transcriptional changes in CTNS-deficient rat kidneys. (**A**) Volcano plot of differently expressed genes (DEGs) in dapagliflozin- versus vehicle-treated *Ctns*^KO^ kidneys (|log2FC| > 0.5, P < 0.05). Genes that are not significantly changed (false discovery rate (FDR) > 0.05) are shown in grey, whereas significantly upregulated and downregulated genes are shown in red and blue, respectively. (**B**) Top 5 enriched biological processes among upregulated and downregulated DEG sets. (**C**) Heatmaps of DEGs in the indicated pathways. (**D**) RT–qPCR validation of selected targets in whole-kidney lysates (relative to vehicle-treated *Ctns*^KO^; n = 4 rats per condition). Statistical analyses were calculated by an unpaired two-tailed Student’s t-test in (**D**). *p < 0.05 and **p < 0.01 versus vehicle-treated wild-type or vehicle-treated *Ctns*^KO^ rats.

Dapagliflozin partially reversed transcriptional alterations in fatty acid metabolism, mitochondrial homeostasis, glucose handling, and ketone-body metabolism (**Figure 6C and 6D**). Circulating ketone bodies increased modestly (**Table S2**), consistent with systemic substrate utilization. Pathway enrichment also highlighted PPARα signalling (**Figure 7A**).^26^ PPARα protein increased (**Figure 7B**), lipid droplet content decreased (Perilipin2^+^; **Figure 7C**), and PT differentiation/transport markers (*Slc6a19, Slc13a3, Slc5a12, Slc27a2*) correlated with lipid metabolic regulators, including the PPARα target *Acox1 and* PPARα coactivator *Ppargc1a* (**Figure 7D-7E<u>; Figure S6A and S6B</u>**; ref.^27^). Moreover, *Acox1*, *Ppargc1a*, and *Lrp2* expression inversely correlated with urinary albumin excretion (**Figure 7F**), linking metabolic transcriptional signatures to improved tubular function.

**Figure 7.**
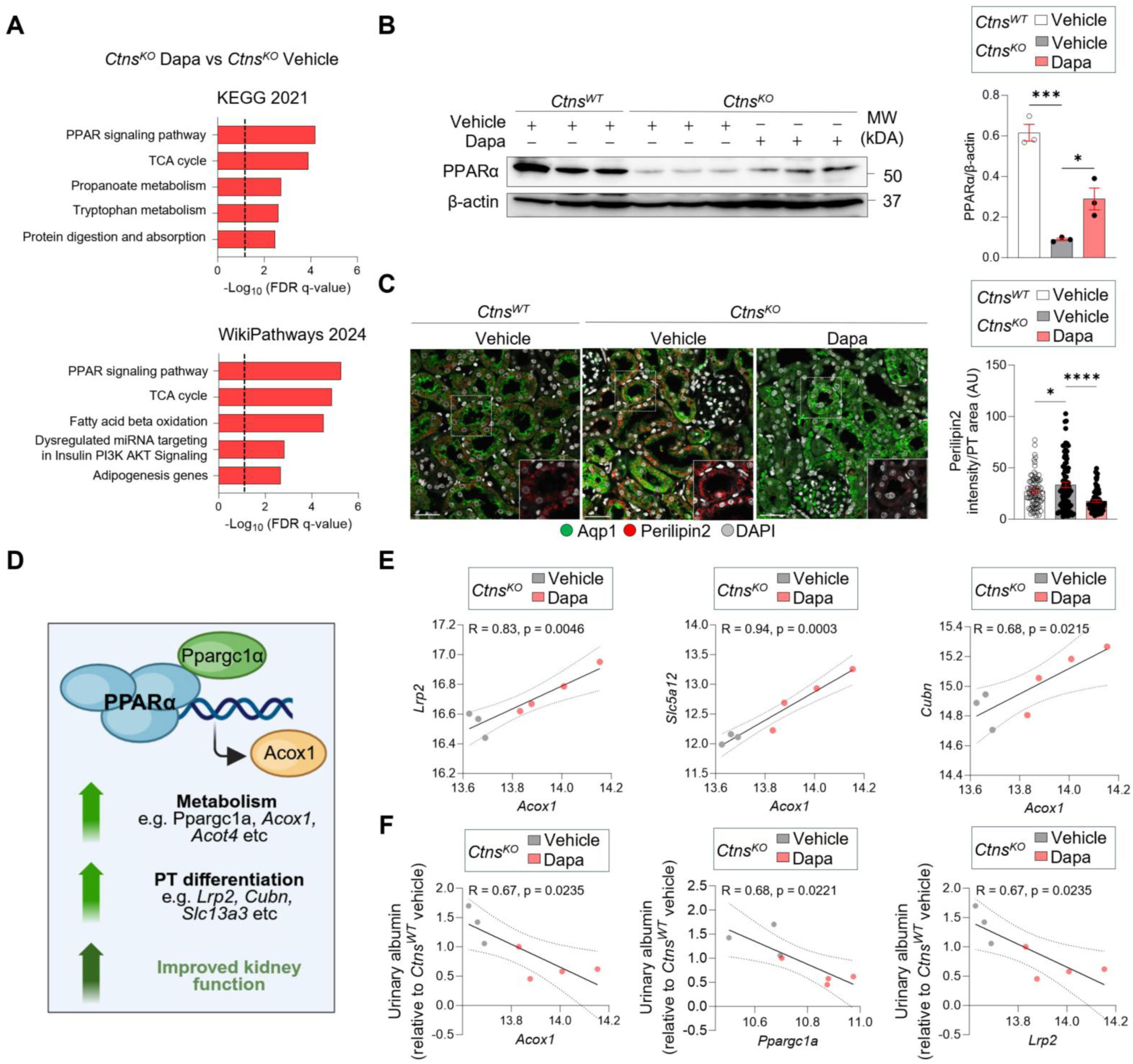
**Dapagliflozin is associated with activation of PPAR**_α_**-linked lipid metabolic programs in CTNS-deficient kidneys.** (**A**) Top 5 upregulated pathways identified across the indicated databases (Gene Ontologies – Biological Processes). (**B**) Immunoblotting and quantification of PPARα in whole-kidney lysates after 4 weeks of treatment (n = 3 animals per group). (**C**) Representative confocal micrographs and quantification of Perilipin2 (red) in the kidney PTs after 4 weeks of treatment (n = 3 fields pooled from n = 3 biologically independent samples). (**D**) Pearson correlation between the expression of the indicated genes and *Acox1* in *Ctns^KO^* kidneys. Plots show normalized Log_2_ expression values (n ≥ 3 animals per condition). (**E**) Pearson correlation between urinary albumin levels and the expression of the indicated genes in *Ctns^KO^* kidneys. Plots show relative albumin levels and normalized Log_2_ expression values (n ≥ 3 animals per condition). Data are mean ± SEM. Statistical analyses were performed using one-way ANOVA followed by Sidak’s multiple comparisons test in (**B** and **C**). For correlations (**E** and **F**), statistics were assessed by Pearson correlation (two-tailed). Dotted white boxes indicate images at higher magnification. Nuclei counterstained with DAPI (grey). Scale bars, 50µm.

Together, these data indicate that SGLT2 inhibition is associated with reduced lysosome-associated mTORC1 activity and induction of PPARα-linked metabolic programs, coinciding with improved PT differentiation and function downstream of CTNS deficiency and cystine storage.

## Discussion

Cystinosis, caused by loss-of-function mutations in CTNS, is characterized by lysosomal cystine accumulation, impaired receptor-mediated endocytosis, and early-onset proximal tubulopathy that progresses to kidney failure.^3,4^ Here, we show across zebrafish, rodent, and PT cell models that SGLT2 inhibition improves proximal tubular phenotypes downstream of cystine storage, without reducing tissue cystine burden, supporting a therapeutic approach that targets pathogenic signalling rather than cystine clearance.

Our data reinforce lysosome-associated mTORC1 dysregulation as a central downstream consequence of CTNS deficiency, linking lysosomal storage to loss of PT differentiation and function.^8–10^ Across model systems, SGLT2 inhibition reduced pathological mTORC1 signalling toward wild-type levels and was accompanied by changes consistent with altered organization of the lysosomal nutrient-sensing apparatus (v-ATPase-Ragulator-Rag components). In contrast to direct mTOR pathway inhibitors, which can cause profound pathway suppression and dose-limiting toxicity, SGLT2 inhibition was associated with restoration of autophagy-lysosome readouts and recovery of PT epithelial features, including polarity, receptor expression, and endocytic function, together with reduced proteinuria and improved injury-associated markers.

At the metabolic level, SGLT2 inhibition shifted kidney transcriptional programs toward oxidative metabolism, including pathways related to fatty acid β-oxidation and PPARα signalling, which support PT energy homeostasis and differentiation.^26^ These transcriptional changes coincided with improved PT identity and function as assessed by PT marker restoration and functional endocytic assays, linking metabolic rewiring to tubular recovery. SGLT2 inhibition is also known to engage systemic metabolic adaptations.

In our study, circulating ketone bodies increased modestly, consistent with altered substrate utilization. These changes are best interpreted as a context of pathway engagement rather than a primary driver of rescue, particularly given the cellular evidence that PT differentiation and lysosome/autophagy phenotypes improve downstream of CTNS loss. Human SLC5A2 loss-of-function provides a genetic framework for these findings, supporting the concept that reduced PT glucose flux can be accommodated long-term by metabolic adaptation rather than intrinsic kidney injury.

Collectively, these results integrate with broader evidence that SGLT2 inhibitors confer kidney protection beyond glucose lowering across diabetic and non-diabetic kidney diseases.^27–29^ By reducing proximal tubular workload and reshaping metabolic state, SGLT2 inhibition has been linked to improved mitochondrial homeostasis and dampened inflammatory/fibrotic responses.^30,31^ Our findings extend this paradigm by indicating that SGLT2 inhibition can mitigate lysosome-driven signalling defects in cystinosis, suggesting a potential strategy for inherited proximal tubulopathies and other kidney disorders with maladaptive mTORC1 activation.^32–35^

From a translational perspective, SGLT2 inhibitors are orally available, widely prescribed, and approved for pediatric use (≥10 years) in type 2 diabetes.24 Acting downstream of cystine storage, SGLT2 inhibition could complement cysteamine, with potential additive benefit. Early initiation — guided by sensitive biomarkers such as urinary LMW proteins — may maximize efficacy before irreversible structural damage. Prospective studies in cystinosis will be needed to define durability and safety, with attention to volume status, electrolyte balance, and rare but recognized risks such as ketosis in susceptible settings.

In summary, SGLT2 inhibition emerges as a mechanistically grounded and clinically actionable strategy that improves PT differentiation and function in cystinosis while lowering pathological mTORC1 activity and engaging oxidative metabolic programs. These findings broaden the therapeutic landscape for cystinosis and support repurposing SGLT2 inhibitors for mTORC1-driven proximal tubule disorders.

## Limitations of the study

Several mechanistic questions remain. First, while our data are consistent with modulation of lysosome-centered nutrient sensing, we did not test whether enforced retention of Ragulator-Rag complexes at lysosomes is sufficient to attenuate SGLT2i-mediated rescue. Second, the contribution of energy-sensing pathways (e.g., AMPK and SIRT1) and their integration with PPARα-linked transcriptional programs remains to be defined. Third, whether SGLT2 inhibition supports durable re-establishment of PT identity (e.g., via epigenetic remodelling) and whether established dedifferentiation and fibrosis can be reversed at later disease stages warrants further study. Finally, systemic contributors—including hemodynamic effects, hormonal signalling, and gut-kidney metabolic crosstalk^36^ — may shape treatment responses *in vivo* and should be addressed in future work. Despite these limitations, the cross-species consistency of the phenotypic and signalling rescue supports a consistent therapeutic effect and provides a framework for follow-up studies to define pathway hierarchy and translational boundaries.

## Materials and Methods

### Antibodies and reagents

The following antibodies were used in this study: anti-LAMP1 (Santa Cruz Biotechnology, sc-19992, 1:1000); anti-LC3 (PM036, MBL, 1:200); anti-p62/SQSTM1 (ab56416, Abcam, 1:500); anti-p62/SQSTM1 (ab109012, Abcam, 1:500), anti-cathepsin D (69854, Cell Signaling Technology; 1:500), anti-phospho-S6 Ribosomal Protein (Ser235/236; 4858, Cell Signaling technology; 1:500), anti-S6 Ribosomal Protein (2217, Cell Signaling Technology, 1:500), anti-phosphoLJ4E-BP1 (Ser65; 9451, Cell Signaling Technology; 1:500), anti-4E-BP1 (9644, Cell Signaling Technology; 1:500), Perlilipin2 (ab108323, Abcam; 1:400), anti−RagC (5466, Cell Signaling Technology; 1:500); anti-Lamptor (8975, Cell Signaling Technology; 1:500), anti-ATP6v1b2 (14617, Cell Signaling Technology; 1:500), anti-HA (11867423001, Roche; 1:1000), anti-acetylated-tubulin (T7451, Sigma-Aldrich; 1:500), anti-Aqp1(ab9566, Abcam; 1:400), anti-Aqp1 (AB2219, EMD Millipore Corp; 1:400), anti-PCNA (M0879, Dako A/S; 1:500), anit-SGLT2 (85626, Abcam; 1:500), anti-SGLT2 (28683, Thermo Fischer Scientific; 1:500), anti-Caspase 3 (9661, Cell Signaling Technology; 1:500), anti-cleaved caspase 3 (9662, Cell Signaling Technology; 1:500), anti-β-actin (A5441, Sigma-Aldrich; 1:1000), anti-γ-tubulin (clone GTU-88, Sigma-Aldrich; 1:500), anti-Transferrin (A0061, Dako A/S; 1:500), anti-Vdbp (A0021, Dako A/S; 1:500), anti-Vimentin (Ab92547, Abcam; 1:400), anti-Lrp2 (1:1000) was kindly provided by P. Verroust and R. Kozyraki (INSERM, Paris, France), donkey-anti rabbit IgG, cross-absorbed secondary antibody, Alexa Fluor 488 (A10040, Thermo Fischer Scientific; 1:400), donkey-anti mouse IgG, cross-absorbed secondary antibody, Alexa Fluor 488 (A21202, Thermo Fischer Scientific; 1:400), donkey-anti rat IgG, cross-absorbed secondary antibody, Alexa Fluor 488 (A21202, Thermo Fischer Scientific; 1:400), donkey-anti rat IgG, cross-absorbed secondary antibody, Alexa Fluor 647 (A31573, Thermo Fischer Scientific; 1:400), donkey-anti goat IgG, cross-absorbed secondary antibody, Alexa Fluor 633 (A21094, Thermo Fischer Scientific; 1:400), donkey-anti goat IgG, cross-absorbed secondary antibody, Alexa Fluor 546 (A11056, Thermo Fischer Scientific; 1:400), donkey-anti rat IgG, cross-absorbed secondary antibody, Alexa Fluor 546 (A11081, Thermo Fischer Scientific; 1:400).

Compounds included Dapagliflozin (BMS-512148, MedChemExpress), Empagliflozin (BI-10773, MedChemExpress), ATP-competitive mTOR inhibitor Torin1 (CAS 1222998-36-8, TOCRIS Bioscience), allosteric mTOR inhibitor rapamycin (R0395-1MG, Sigma-Aldrich), Cysteamine (M6500, Sigma-Aldrich). The cells used in this study were negatively tested for mycoplasma contamination using MycoAlert™ Mycoplasma Detection Kit (LT07-118, Lonza).

### Experimental model and subject detail

**Ethical compliance.** All the animal experiments were approved by the Cantonal Veterinary Office Zurich (license no. ZH230/2019 and no. ZH139/2022) and performed in accordance with the guidelines of the University of Zurich.

**Zebrafish model.** The *ctns* zebrafish model was generated and validated as previously described.^8,9,22^ For functional studies of PT endocytosis, a stable transgenic zebrafish line expressing mCherry-tagged half vitamin D binding protein (½vdbp-mCherry) under a liver-specific promoter was crossed with *ctns* zebrafish.^8,9,22^ Zebrafish were maintained under standard conditions at 28°C on a 14h light/10h dark cycle at 28°C. Larvae were fed a standard diet (Zebrafeed 100–200, SPAROS) starting at 5 days post-fertilization (dpf). All experiments were performed using larvae between 5 and 14dpf. Sex determination was not performed, as sex cannot be reliably assigned at these developmental stages. For pharmacological treatments, zebrafish larvae were exposed from 5dpf to system water containing DMSO, dapagliflozin (100-300nM), empagliflozin (100-1000nM), cysteamine (0.5mM) or rapamycin (200nM) for 9 consecutive days.

**Rodent models.** All animal experiments were performed using age- and sex-matched *Ctns* knockout mice (C57BL/6 background) and rats (Sprague-Dawley background), together with their respective wild-type littermate controls. Mice aged 24 weeks were used for primary cell isolation and mechanistic studies. Rats were used for *vivo* therapeutic validation experiments. Animals were housed under specific pathogen-free conditions in temperature-controlled (22–25°C) and humidity-controlled (50–60%) facilities with a 12hour light/12h dark cycle. Standard chow and water were provided *ad libitum*. Animals were monitored regularly for general health and body weight throughout the experimental period. For pharmacological studies, 10-week-old female *Ctns* knockout rats were provided with drinking water containing dapagliflozin (10 mg kg⁻¹ day⁻¹) for 4 weeks. Vehicle-treated control animals received standard drinking water. Drug dosage was adjusted based on average daily water intake. At the end of the treatment period, rats were euthanized, and kidneys were harvested for biochemical, histological, and molecular analyses. All procedures were conducted in accordance with institutional and national guidelines for animal care and were approved by the relevant animal ethics committees.

**Isolation of PT segments from mouse kidneys and primary cell culture.** Primary mouse proximal tubule cells were isolated from kidneys from *Ctns* knockout mice and their wild-type littermate controls for the isolation of primary mouse proximal tubule cells (mPTCs). Kidneys were harvested immediately after sacrifice, and PT segments were freshly microdissected, as previously described.^8,9^ Isolated PT segments were seeded onto collagen-coated chamber slides (C7182, Sigma-Aldrich) or collagen-coated 6- or 24-well plates (145380 or 142475, Thermo Fisher Scientific). Cells were cultured at 37°C in a humidified incubator with 5% CO₂ in DMEM/F12 medium (21041-025, Thermo Fisher Scientific), supplemented with 5% dialyzed fetal bovine serum (FBS), 15mM HEPES (H0887, Sigma-Aldrich), 0.55mM sodium pyruvate (P2256, Sigma-Aldrich), and 0.1mLLJL⁻¹ non-essential amino acids (M7145, Sigma-Aldrich). Additional supplements were provided using the SingleQuots® kit (CC-4127, Lonza) and included hydrocortisone, human epidermal growth factor (EGF), epinephrine, insulin, triiodothyronine, transferrin, and gentamicin/amphotericin. Culture medium was adjusted to pH 7.40 and an osmolality of 325mOsm/kg and was replaced every 48 hours. After 6-7 days, confluent epithelial monolayers emerged from the tubular fragments and were characterized by high receptor-mediated endocytic activity. All cultures were routinely tested and confirmed negative for mycoplasma contamination.

### Method details

**Quantification of urine ½vdbp-mCherry in zebrafish.** Urinary excretion of low-molecular-weight proteins was quantified using an ELISA-based assay for ½vdbp–mCherry. Zebrafish larvae were individually placed in a 48-well plate, with each well containing one larva in 500μL of zebrafish facility system water. Plates were maintained at 28°C for 16 hours to allow accumulation of excreted urine. Following incubation, the conditioned water from each well was collected and used directly for analysis. Levels of ½vdbp–mCherry were quantified using a mCherry ELISA kit (ab221829, Abcam).

**Kidney function and metabolic parameters**. For evaluation of kidney function and metabolic parameters, rats were housed individually in metabolic cages overnight with free access to food and water. Urine was collected on ice, and physiological parameters including body weight, water intake, and urine output (diuresis), were recorded. Blood samples were collected from the sublingual vein under anesthesia using isoflurane. Urine and blood biochemical parameters were quantified using the UniCel DxC 800 PRO Synchron Clinical System (Beckman Coulter). Urinary albumin and LMW proteins were assessed by Coomassie Blue staining following SDS-PAGE. Protein loading was normalized to total urinary protein^37^, which was determined using ProtoBlue Safe reagent (EC-722, National Diagnostics). Ketone body plasma samples were measured after the treatment using the Ketone Body Assay Kit (ab 318308, Abcam). The assays for acetoacetate (AcAc) and β-hydroxybutyrate (BHB) were performed simultaneously and in duplicates. The total ketone body concentration was calculated as [Total ketone bodies] = [AcAc] + [BHB].

**Cellular treatments.** Primary mPTCs were treated with either dapagliflozin (2µM, unless otherwise indicated) or with the ATP-competitive mTOR inhibitor Torin1 (250nM). Treatments were performed in complete cell culture medium supplemented with 0.1% dialyzed FBS for 16 hours. Vehicle-treated cells received the corresponding concentration of solvent. Following treatment, cells were processed for downstream analyses as described in the respective methods sections.

**Adenovirus transduction.** For lysosomal expression studies, an adenoviral construct encoding 3×HA-tagged TMEM192 was used at a fixed volume of viral stock per well (0.2µL per 250’000 cells). Recombinant adenoviral particles were obtained from Vector Biolabs (University City Science Center, Philadelphia, USA). Primary mPTCs were seeded onto collagen-coated 6-well tissue culture plates. Primary mPTCs were seeded onto collagen-coated 6-well tissue culture plates and allowed to attach for 24 hours until reaching approximately 70-80% confluence. Adenoviral transduction was performed by incubating cells with viral particles for 6 hours at 37°C in complete culture medium. Following transduction, cells were washed and maintained in fresh medium, which was replaced every 48 hours. Cells were harvested 48 hours after transduction for downstream expression and biochemical analyses.

**Lysosome immunoprecipitation.** Lysosomes were isolated by immunoprecipitation from cells transiently expressing 3×HA-tagged TMEM192, as previously described.^8^ Cells were used in collagen-coated 6-well plates and cultured to near confluency within 24 hours.

Following removal of culture medium, cells were scraped into 200µL of ice-cold fractionation buffer containing 50mM KCl, 90mM K-glutamate, 1mM EGTA, 5LJmM MgCl_2_, 50mM sucrose, 5mM glucose, 20mM HEPES (pH 7.4), supplemented with protease inhibitors (1836153001, Roche) and phosphatase inhibitors (PhosSTOP, 04906845001, Sigma Aldrich). Cell suspensions were collected by centrifugation (200LJ×LJg, 5 min, 4°C). Pellets were resuspended in 1mL of fresh fractionation buffer, and approximately 4LJ×LJ10LJ cells were mechanically disrupted by passing the suspension 10 times through a 23G needle attached to a 1mL syringe. Lysates were centrifuged (400LJ×LJg, 10 min, 4LJ°C) to remove nuclei and debris, yielding a post-nuclear supernatant. Protein concentration was determined using the Bradford assay (23246, Thermo Fisher Scientific). Equal amounts of protein were incubated with 50μL of anti-HA conjugated magnetic Dynabeads (88837, Thermo Fisher Scientific) for 1 hour at 4°C with gentle end-over-end rotation. Beads were then washed four times with fractionation buffer to remove unbound material. For immunoblotting analysis, the lysosome-bound beads were resuspended in 40μL of Laemmli sample buffer, boiled at 95°C for 10 minutes, resolved by 12% SDS-PAGE, and analyzed by immunoblotting.

**Endocytic uptake assay in mPTCs.** Endocytic activity in mPTCs was assessed using fluorescently labelled bovine serum albumin (BSA). To allow ligand binding, cells were incubated with 50μg/mL BSA–Alexa Fluor 488 (A34785, Thermo Fisher Scientific) in HEPES-buffered Dulbecco’s Modified Eagle’s Medium (DMEM) for 60 minutes at 4°C. Following ligand binding, cells were shifted to 37°C in complete growth medium and incubated for 20 minutes to permit endocytic internalization. Cells were subsequently fixed and processed for immunofluorescence analysis as described below.

**Lysosomal activity assay.** Lysosomal proteolytic activity in primary mPTCs was assessed using Bodipy-FL-Pepstatin A (P12271, Thermo Fischer Scientific), a fluorescent probe that binds active cathepsin D. Cells were incubated with 1µM BODIPY FL–Pepstatin A in Live Cell Imaging medium for 1 hour at 37°C, followed by fixation and analysis by confocal microscopy. Quantification of lysosomal activity was performed by measuring the number of Pepstatin A-positive structures per cell were quantified using the open-source cell image analysis software CellProfiler^TM^.^41^

**Cell viability assay.** Cell viability of mPTCs following pharmacological treatments was assessed using the MTT assay (ab211091, Abcam), according to the manufacturer’s protocol. Briefly, cells were washed three times with PBS and incubated with MTT solution (0.5mg/mL) prepared in complete culture medium. After incubation for 4 hours at 37°C, during which intracellular formazan crystals formed, the medium was removed and crystals were solubilized using dimethyl sulfoxide (DMSO; 276855, Sigma-Aldrich). Absorbance was measured at 570nm using a microplate reader. Cell viability was expressed relative to vehicle-treated control conditions.

**Cell proliferation**. Proliferation of mPTCs was assessed using the Click-iT™ Plus EdU Alexa Fluor™ 488 Imaging Kit (C10637, Life Technologies). Cells were incubated with EdU for 16 hours at 37°C to allow incorporation during DNA synthesis. Following fixation and permeabilization, incorporated EdU was detected via the Click-iT reaction. The percentage of EdU-positive nuclei was quantified per field. Nuclei were counterstained with DAPI. Images were acquired using a Leica SP8 confocal laser scanning microscope.

**Cystine measurements**. Cystine levels were quantified in rat kidney tissue and zebrafish larvae as previously described.^8^ All values were normalized over total protein concentration.

**RNA Extraction, cDNA Synthesis, and RT-qPCR Analysis.** Total RNA was isolated from rat and zebrafish tissues using the Aurum™ Total RNA Fatty and Fibrous Tissue Kit (732-6830, Bio-Rad) with DNase I treatment, and from primary mPTCs using the RNAqueous® Kit (AM1912, Ambion). One microgram of RNA was reverse transcribed using the iScript™ cDNA Synthesis Kit (170-8890, Bio-Rad). Quantitative real-time PCR (RT-qPCR) was performed using the CFX96™ Real-Time PCR Detection System (Bio-Rad) and iQ™ SYBR Green Supermix (170-8880, Bio-Rad). Reactions were run in duplicates using gene-specific primers (100nM; sequences in Table S3). Cycling conditions were at 95°C for 3 minutes, followed by 40 cycles at 95°C for 15 seconds and at 60°C for 30 seconds. Primer specificity was verified by melt-curve analysis, and selected PCR products were sequence-verified using the BigDye™ Terminator v3.1 Cycle Sequencing Kit (4337455, Applied Biosystems) on an ABI 3100 capillary sequencer. Primer efficiencies were determined using standard dilution curves. Relative changes in the expression of target genes were calculated using the 2−ΔΔCt method, with *eef1a1a*, *Hrpt,* or *Gapdh* as the reference gene.

### Bulk RNA sequencing

*Sample preparation*. Total RNA was extracted from whole kidney samples of *Ctns* rat models using the RNeasy Plus Mini Kit (74134, Qiagen). RNA quantity and integrity were assessed before library preparation. RNA-seq libraries were generated from 100ng of total RNA per sample using the TruSeq Stranded mRNA Library Prep Kit (Illumina). The quality of both total RNA and prepared libraries was assessed using a Fragment Analyzer (Agilent).

Paired-end sequencing (2×100bp) was carried out on a NovaSeq 6000 platform (Illumina) *Bioinformatics analysis*. Raw sequencing data were processed and analyzed using established RNA-sequencing workflows. Principal component analysis (PCA), generation of heat maps and overrepresentation analysis (ORA) were performed using Network Analyst 3.0.^38^ Data were log_2_-normalized and analyzed using the DESeq2 algorithm.^39^ Volcano plots and ORA scatter plots were generated using the exploreDE Shiny app.^40^ Genes were considered differentially expressed based on the thresholds indicated in the corresponding figure legends.

**Immunofluorescence Staining and Confocal Microscopy.** *Rat and mouse tissue slides*. Fresh rat and mouse kidneys were fixed by transcardial perfusion with 4% PFA, dehydrated, paraffin-embedded and sectioned at 5μm. Sections were deparaffinized in xylene (534056, Sigma-Aldrich), rehydrated through a graded ethanol series, and subjected to antigen retrieval in sodium citrate buffer (pH 6.0). Sections were then quenched with 50mM NH_4_Cl, blocked in 3% BSA prepared in PBS containing calcium and magnesium (PBS Ca/Mg; D1283, Sigma-Aldrich), and incubated overnight at 4°C with primary antibodies. After washing in PBS containing 0.1% Tween 20, sections were incubated with Alexa fluorophore-conjugated secondary antibodies (Invitrogen), counterstained with DAPI (1µM; D1306, Thermo Fisher Scientific), mounted using ProLong® Gold Antifade Reagent (P36930, Thermo Fisher Scientific) and imaged by confocal microscopy.

*Cultured mPTCs.* Cells were fixed with 4% PFA for 10 minutes, quenched with 50mM NH_4_Cl, permeabilized for 20 minutes in blocking buffer containing 0.1% Triton X-100 and 0.5% BSA in PBS, and incubated overnight at 4°C with primary antibodies. Cells were washed repeatedly with PBS and incubated with fluorophore-conjugated Alexa secondary antibodies (Invitrogen). Nuclei were counterstained with 1μM DAPI, and coverslips were mounted using ProLong® Gold Antifade Reagent (P36930, Thermo Fisher Scientific).

*Image processing and quantification*. Images were acquired using a Leica SP8 confocal laser scanning microscope (Center for Microscopy and Image Analysis, University of Zurich), with a Leica APO × 63 NA 1.4 oil-immersion objective under constant acquisition settings.

Images were collected at a resolution of 1024LJ×LJ1024 pixels, with 8-16-line scan averages, and the pinhole set to 1 Airy unit for each emission channel to ensure linear signal detection (intensity range: 1–254). Quantitative image analysis was performed on 3 randomly selected fields per section, each containing at least 30 PTs, identified by Aqp1-positive structures for the rat tissue. For fixed mPTCs, 3 randomly selected fields per chambre were analysed, each containing a minimum of 20 cells. Quantitative image analysis was performed using the open-source image analysis software CellProfiler^TM,41^ and Fiji (ImageJ, NIH).

**Imaging of zebrafish larvae.** Quantitative analysis of fluorescent signals was performed using one entire pronephric tubule per larva, with all imaging parameters kept constant across samples. High-resolution imaging of fluorescent vesicles in proximal tubules was carried out using a multi-photon fluorescence microscope (Leica SP8 MP DIVE Falcon,

Leica Microsystems, Center for Microscopy and Image Analysis, University of Zurich) equipped with a ×25/1.0 NA water immersion objective (HC IRAPO L, Leica Microsystems). For the detection of mCherry fluorescence, excitation was achieved using a tunable laser at a wavelength of 1040nm. Acquired z-stack images were deconvolved using Huygens software (Scientific Volume Imaging) and subsequently segmented with the Imaris software (Oxford Instruments). Quantification of fluorescence intensity per z-stack was performed using Fiji (ImageJ, NIH).

**Quantification of kidney fibrosis.** Renal fibrotic tissue was visualized on paraffin-embedded kidney sections using Picro-Sirius Red solution (ab150681, Abcam), performed according to the manufacturer’s instructions. Following staining, slides were mounted using toluene-based mounting medium (SP15-500, Thermo Fisher Scientific). Whole-slide images were acquired using a Zeiss Axio Scan.Z1 automated slide scanner (Center for Microscopy and Image Analysis, University of Zurich) equipped with a Plan Apochromat ×40 NA 0.95 air-immersion objective. Quantification of fibrosis was performed using Fiji software (ImageJ, NIH) by color deconvolution of Sirius Red-stained collagen (red) and non-collagen components (orange). The fibrotic area was expressed as a percentage of the total tissue surface, as previously described by Bienaimé et al.^42^

**Immunoblotting.** Proteins were extracted from animal tissues or cultured cells using lysis buffer supplemented with protease inhibitors (1836153001, Roche) and phosphatase inhibitors (PhosSTOP, 04906845001, Sigma Aldrich), followed by brief sonication and centrifugation (16,000LJ×LJg, 10min, 4°C). Protein concentrations were determined using standard colorimetric assays. Equal amounts of protein (20μg) were denatured under reducing conditions, separated by SDS-PAGE and transferred to PVDF membranes, which were blocked with 5% non-fat milk in tris-buffered saline containing 0.1% Tween-20.

Membranes were incubated overnight at 4°C with the primary antibodies, followed by incubation with HRP-conjugated secondary antibodies. Proteins were detected using enhanced chemiluminescence (ECL; WBKLS0050, Millipore, Life Technologies) and imaged with a ChemiDoc MP Imaging System (Bio-Rad Laboratories). Densitometric analysis was performed using Fiji (ImageJ, NIH), and signal intensities were normalized to indicated loading controls.

**Quantification and statistical analysis**. Quantification methods are described in the Method Details and in the figure legends. Data are presented as the mean ± standard error of the mean (SEM). Statistical analyses were performed using GraphPad Prism software v.10.1.2 (GraphPad Software). Comparisons among multiple experimental groups were conducted using one-way analysis of variance (ANOVA), followed by Tukey’s, Šídák’s, or Dunnett’s multiple-comparison tests, as appropriate. Comparisons between two groups were performed using two-tailed unpaired or paired Student’s *t*-tests, depending on the experimental design. Data normality was assessed using the D’Agostino–Pearson omnibus normality test. For non-normally distributed data, statistical significance was determined using the Kruskal–Wallis test with Dunn’s multiple-comparison correction. Pearson correlation coefficients (*R*) were calculated to assess linear relationships between variables, including gene expression levels and quantitative phenotypic readouts. Correlation analyses were performed using log₂-transformed, normalized RNA-sequencing expression values. Scatter plots were generated with fitted linear regression lines and annotated with Pearson correlation coefficients and corresponding two-tailed *P*-values. Statistical significance, *P*-values, and the statistical tests used are indicated in the figure legends. Unless otherwise stated, experiments were independently repeated two to three times with similar results. Investigators were not blinded to group allocation during the experiments or outcome assessments. Outliers were identified using the ROUT method in GraphPad Prism (Q = 1%) and excluded from statistical analyses.

## Resource availability

### Lead contact

Further information and requests for resources and reagents should be directed to the lead contacts, Alessandro Luciani and Olivier Devuyst.

### Material availability

This study did not generate new reagents.

### Data availability

Primary RNA-sequencing data have been deposited on Gene Expression Omnibus with the dataset identifier GSE316607 (https://www.ncbi.nlm.nih.gov/geo/query/acc.cgi?acc=GSE316607).

## Acknowledgements

We thank Nadine Nägele and Sara Botta for technical help; Peter Lary, Chia-wei Lin and the Functional Genomics Center Zurich for the support with the RNA-sequencing data analysis and the cystine measurements; the Center for Microscopy and Image Analysis at the University of Zurich for providing equipment and confocal microscopy assistance; Guglielmo Schiano and the Functional Genomics Center Zurich for assistance; and Prof. Francesco Emma (OPBG, Rome, Italy) for fruitful discussions. We are grateful to the Cystinosis Research Foundation [Irvine, CA, USA; project grants CRFS-2017-007 (O.D. and A.L.), CRFS-2020-005 (O.D. and A.L.) and CRFS-2023-003 (O.D. and A.L.)], the Swiss National Science Foundation (project grant 310030_215715 to A.L.), the University Research Priority Program of the University of Zurich (URPP) ITINERARE-Innovative Therapies in Rare Diseases (O.D. and A.L.).

## Author contributions

Conceptualization: A.L., O.D.; Supervision: A.L., O.D. Methodology: S.K., A.L., O.D.; Investigation: S.K., Z.C., P.K.; Resources: Z.C.; Data curation: S.K.; Formal analysis: S.K.; Writing – original draft: A.L., O.D. S.K.; Writing – review & editing: A.L., O.D., S.K.

## Disclosure and competing interest statement

The authors declare no competing interests.

## Notes

### Competing Interest Statement

The authors have declared no competing interest.

