## Supplemental information for "SGLT2 Inhibitors Rescue Lysosomal mTORC1 Hyperactivity and Proximal Tubulopathy in Preclinical Models of Cystinosis"

Keller et al.

Figure S1-S6

Table S1-S3

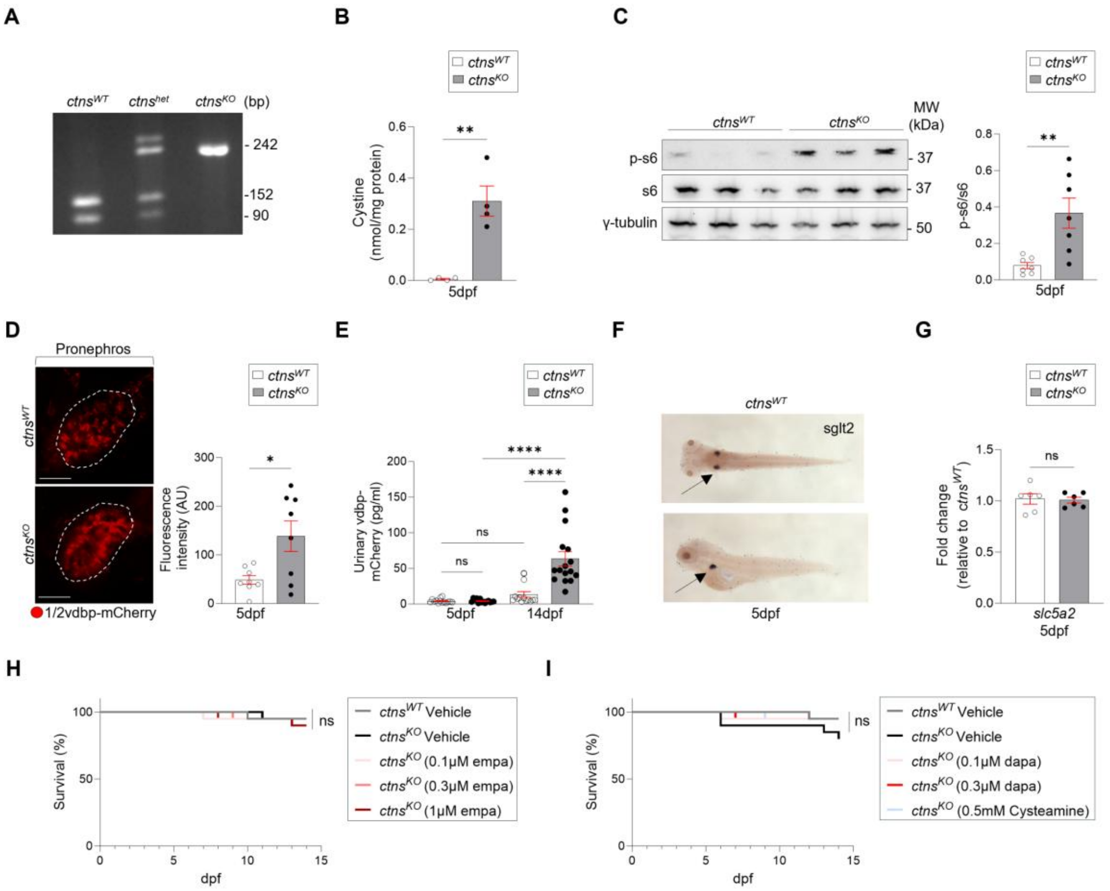


**Figure S1. Zebrafish model of cystinosis recapitulates key disease hallmarks.** (**A**) Genotyping strategy for ctns zebrafish. PCR amplification of ctns exon 3 followed by AciI digestion distinguishes alleles: the WT PCR product is cut into 152bp and 90bp fragments, whereas the mutant allele remains undigested (242bp). Genomic DNA was extracted from the caudal region of WT, heterozygous, and homozygous larvae. (**B**) Cystine levels in larvae at 5dpf. Each dot represents a pool of 20 larvae (n = 4 independent pools per genotype). (**C**) Immunoblot and quantification of pS6/S6 at 5dpf. Each lane represents a pool of 6 larvae (n = 7 independent pools per genotype). (**D**) Multiphoton imaging and quantification of vdbp–mCherry fluorescence intensity in the pronephros (n = 8 larvae per genotype). (**E**) Urinary mCherry by ELISA at 5 and 14dpf (n = 14 larvae per genotype and timepoint). (**F**) Representative slc5a2/sglt2 expression image in zebrafish larvae from ZFIN. (**G**) slc5a2 mRNA levels in ctns larvae. Expression normalized to *eef1a1a* and expressed relative to WT (n = 6 independent samples per genotype). (**H**) Survival of larvae treated with empagliflozin (empa) from 5 to 14dpf (n ≥ 20 larvae per condition). (**I**) Survival of larvae treated with dapagliflozin (dapa) or cysteamine from 5 to 14dpf (n ≥ 20 larvae per condition). Data are mean ± SEM where applicable. For quantitative comparisons in (**B**, **C**, **D**, **E**, **G**), statistical tests were calculated by an unpaired two-tailed Student’s t test. Survival curves in (**H**, **I**) were compared by Mantel–Cox (log-rank) test. NS, non-significant. Scale bars, 20μm.

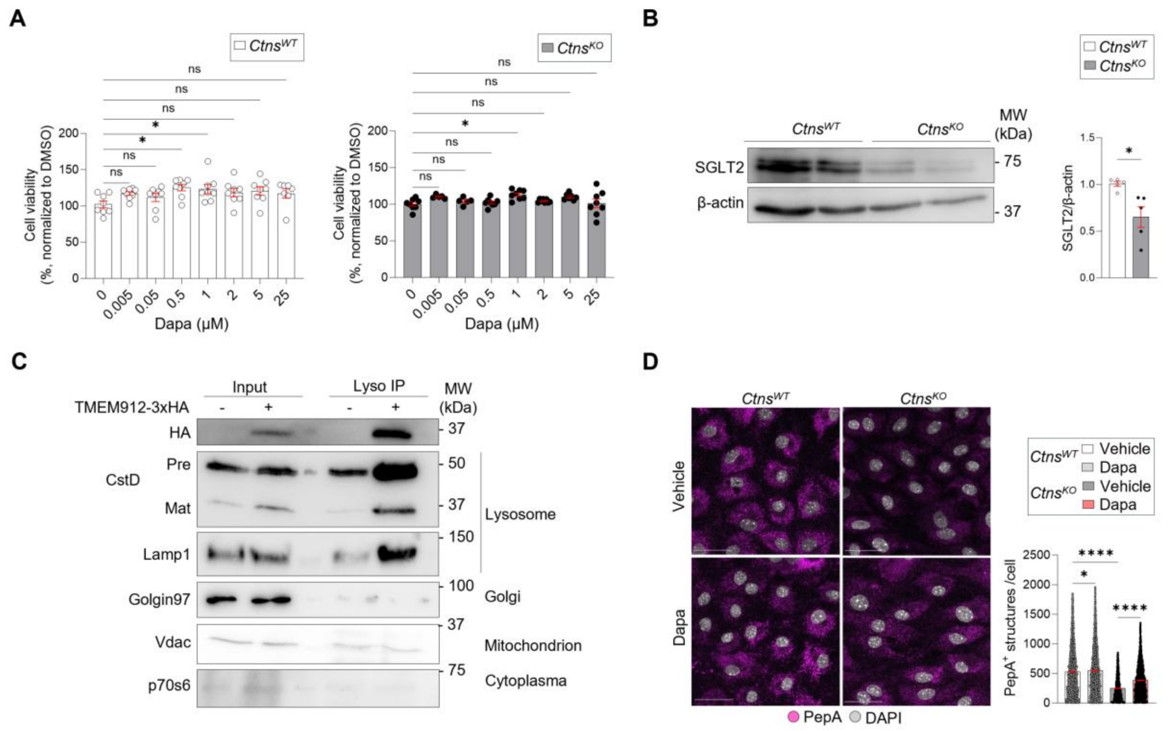


**Figure S2.Toxicity, SGLT2 expression, Lyso-IP validation, and lysosomal activity in mPTCs.** (**A**) mPTCs were treated with dapagliflozin at the indicated concentrations for 16 h. Cell viability was assessed by MTT and normalized to the vehicle control. n = 2 independent mPTCs preparations per condition; 8 technical wells per preparation. (**B**) Immunoblot of SGLT2 and quantification in mPTCs isolated from 24-week-old mice (n = 4 biologically independent samples). (**C**) Cells were loaded with BODIPY FL-Pepstatin A (PepA; 1μM, 1 h, 37°C) and analyzed by confocal microscopy. Quantification of PepA$^+$ structures per cell (n = 3 fields pooled from 3 biologically independent samples). (**D**) Immunoblot validation of lysosome immunopurification (Lyso-IP) from mPTCs transiently expressing TMEM192–3×HA (anti-HA pull-down), showing enrichment of lysosomal markers with depletion of non-lysosomal compartment markers (input vs Lyso-IP). Data are mean ± SEM. Statistical analyses: one-way ANOVA with Sidak's multiple comparisons in (A and **D**) and unpaired two-tailed Student's t test in (**B**). Nuclei counterstained with DAPI (grey). NS, non-significant. Scale bars, 50μm.

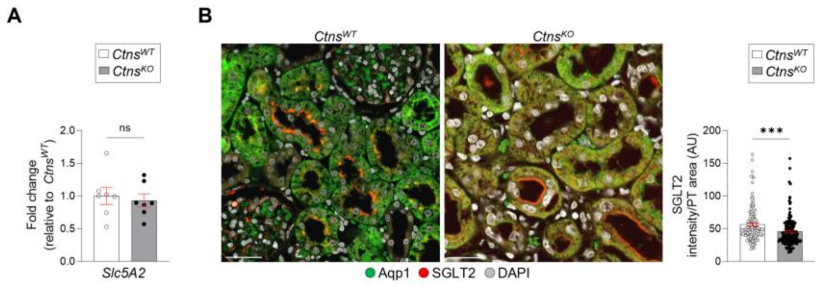


**Figure S3: *Slc5a2*/SGLT2 expression in *Ctns* rat**. (**A**) mRNA levels of *Slc5a2* in the kidney of 12-week-old rats. Gene target expression normalized to *Gapdh* and expression relative to *Ctns*$^{WT}$ (n = 7 biologically independent samples). (**B**) Representative confocal micrographs and quantification of SGLT2i (red) in the kidney PTs of 12-week-old rats (n = 3 fields pooled from n = 3 biologically independent samples per). Statistical analyses were calculated by an unpaired two-tailed Student's t test in (**A** and **B**). NS, non-significant. Scale bars, 50μm.

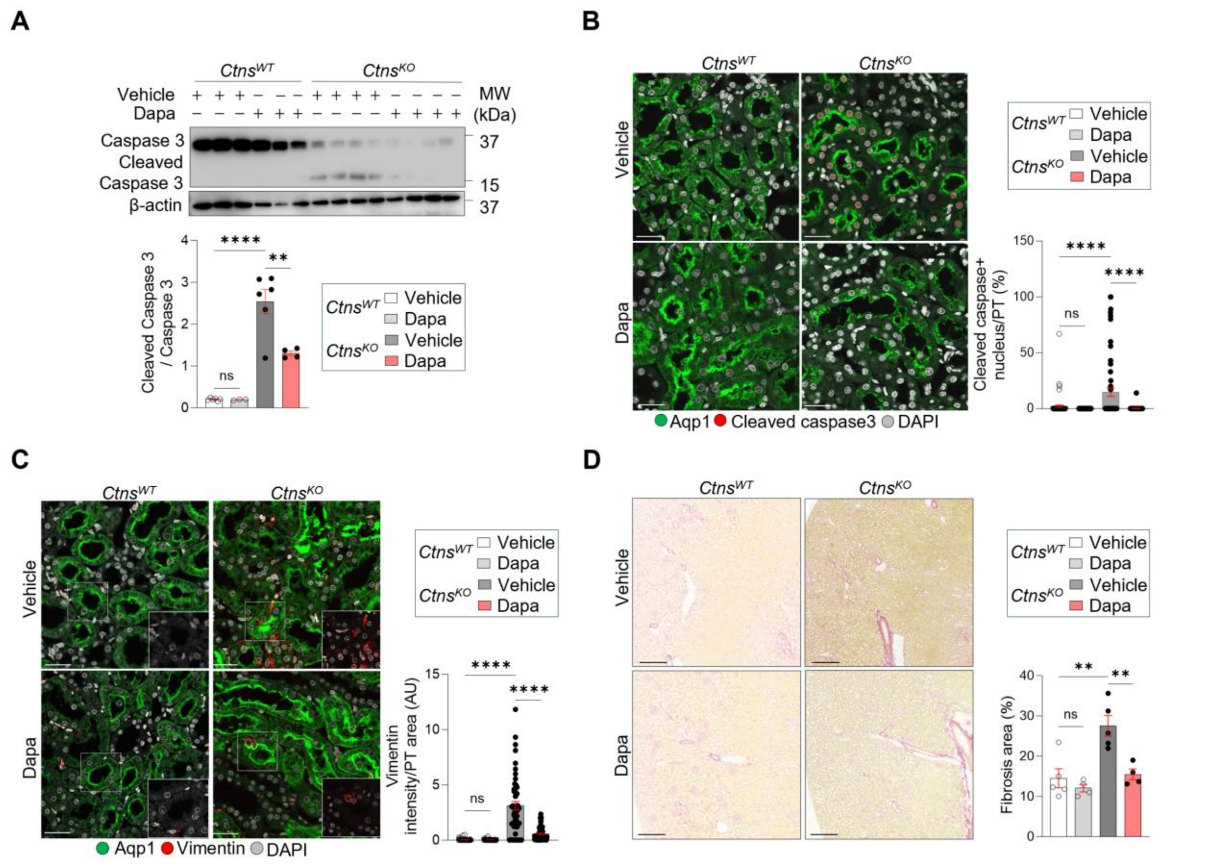


**Figure S4. Dapagliflozin is associated with reduced apoptosis and interstitial fibrosis in CTNS-deficient rats.** (**A**) Immunoblot and quantification of cleaved caspase-3 and total caspase-3 in whole-kidney lysates after 4 weeks of treatment (n ≥ 4 rats per genotype and condition). (**B**) Representative confocal images and quantification of cleaved caspase-3 (red) in proximal tubules after 4 weeks of treatment (nuclei, grey). n = 3 fields pooled from n = 3 biologically independent samples. (**C**) Representative confocal images and quantification of vimentin (red) in proximal tubules after 4 weeks of treatment. n = 3 fields pooled from n = 3 biologically independent samples. (**D**) Picro–Sirius Red staining (representative image) and quantification of fibrotic area fraction in kidney sections after 4 weeks of treatment (n ≥ 4 rats per condition). Data are mean ± SEM. Statistical analyses were calculated by one-way ANOVA followed by Sidak's multiple comparisons test in (**A**-**D**). NS, non-significant. Scale bars, 50μm in (**B** and **C**), 500μm in (**D**).

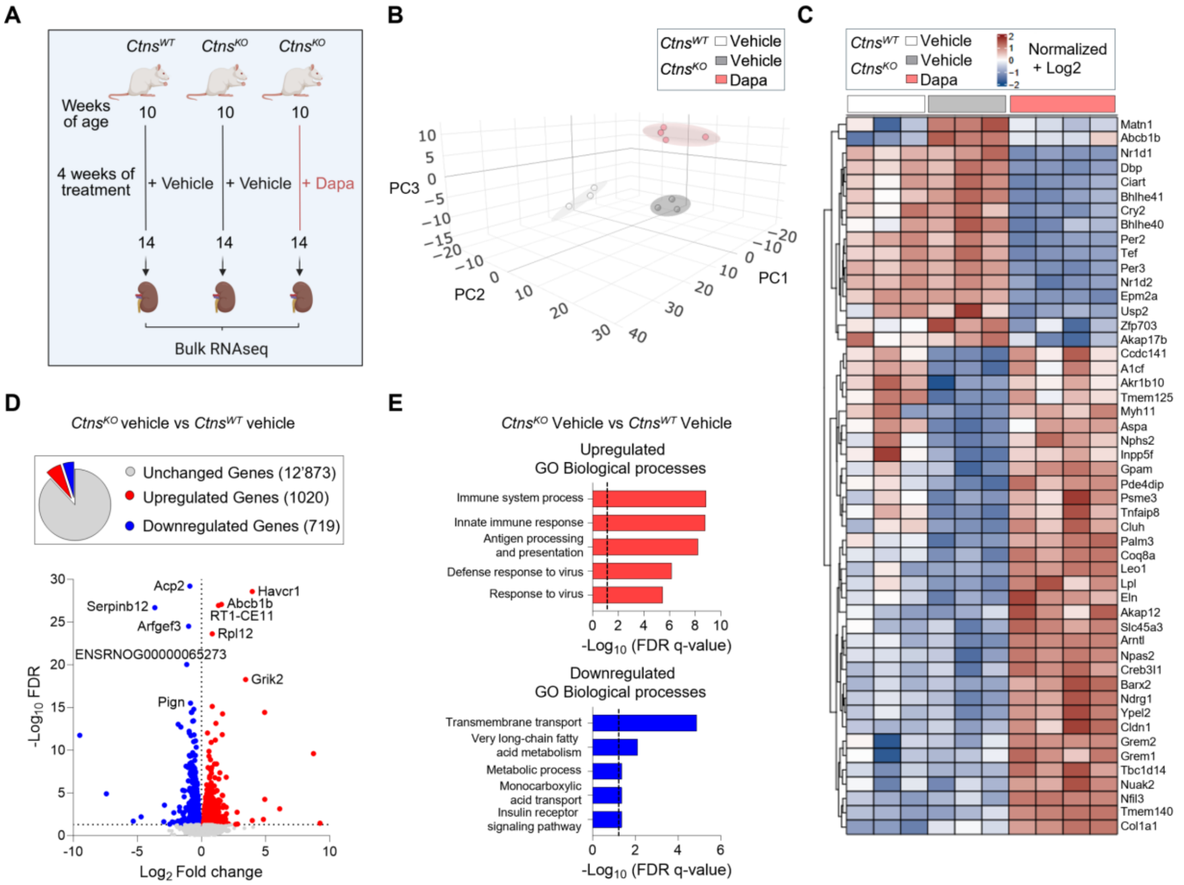


**Figure S5: Bulk RNA sequencing analysis after 4 weeks of dapagliflozin treatment.** (**A**) Study schematic of bulk RNA-seq performed on whole-kidney lysates from *Ctns*$^{WT}$ (vehicle), *Ctns*$^{KO}$ (vehicle), and *Ctns*$^{KO}$ treated with dapagliflozin for 4 weeks. n = ≥ 3 rats per group. (**B**) Principal component analysis (PCA) of kidney transcriptomes across groups. (**C**) Heat map of the top 50 differently differentially expressed genes (DEGs) across the three groups. Genes not significantly changed (false discovery rate (FDR) > 0.05) are shown in grey, whereas significantly upregulated and downregulated genes are shown in red and blue, respectively. (**E**) Top 5 enriched biological processes among upregulated and downregulated DEG sets for *Ctns*$^{KO}$ versus *Ctns*$^{WT}$ rat kidneys.

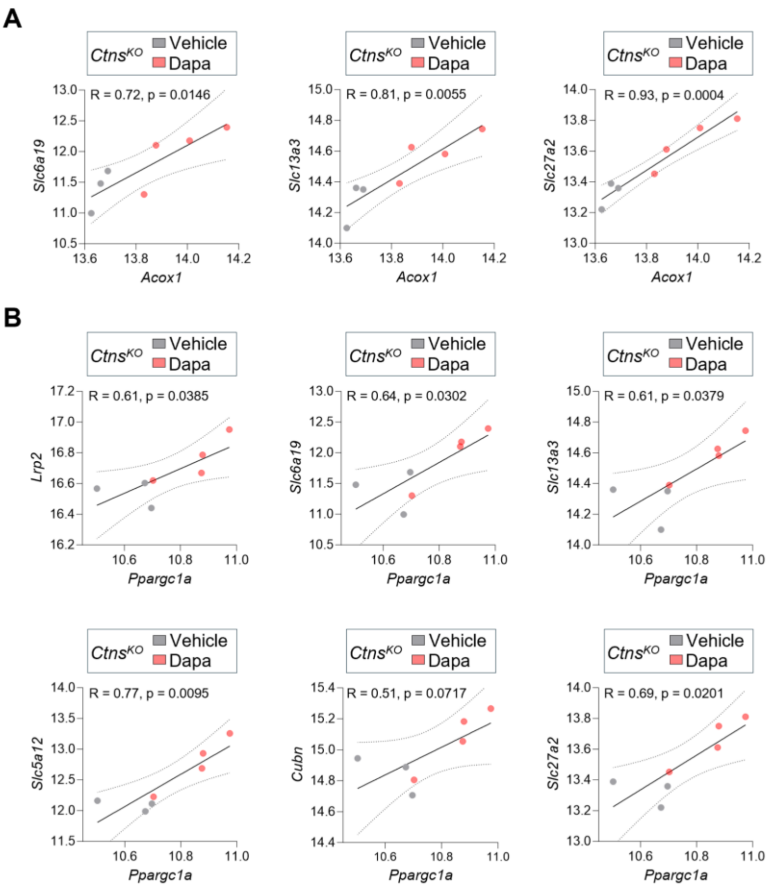


**Figure S6: Correlations between metabolic regulators and proximal tubule differentiation signatures.** (**A**) Pearson correlations between *Acox1* expression and the indicated genes in *Ctns*$^{KO}$ kidneys. (**B**) Pearson correlations between *Ppargc1a* expression and the indicated genes in *Ctns*$^{KO}$ kidneys. Plots show normalized log2 expression values from bulk kidney RNA-seq. Each point represents one animal. Pearson r and two-tailed p values are shown in each panel.

**Table S1: Body weight and urine parameters in *Ctns* rats at baseline and after dapagliflozin treatment**

| Parameter | *Ctns*$^{+/+}$ Baseline Control (n=10) | *Ctns*$^{-/-}$ Baseline Control (n=10) | *Ctns*$^{+/+}$ 2 weeks Control (n=6) | *Ctns*$^{+/+}$ 2 weeks Dapa (n=4) | *Ctns*$^{-/-}$ 2 weeks Control (n=6) | *Ctns*$^{-/-}$ 2 weeks Dapa (n=4) | *Ctns*$^{+/+}$ 4 weeks Control (n=6) | *Ctns*$^{+/+}$ 4 weeks Dapa (n=4) | *Ctns*$^{-/-}$ 4 weeks Contro l (n=6) | *Ctns*$^{-/-}$ 4 weeks Dapa (n=4) |
|---|---|---|---|---|---|---|---|---|---|---|
| Body weight (g) | 245 ± 12.5 | 241 ± 19.5 | 252 ± 9.46 | 243 ± 9.26 | 257 ± 15.5 | 232 ± 12.2$^{\dagger}$ | 259 ± 6.37 | 256 ± 7.53 | 264 ± 14.0 | 238 ± 18.8$^{\dagger}$ |
| Water intake (ml in 16h) | 26.0 ± 6.76 | 23.7 ± 4.52 | 30.6 ± 8.71 | 48.3 ± 5.03** | 26.8 ± 6.24 | 42.5 ± 2.38$^{\dagger\dagger}$ | 32.3 ± 7.20 | 51.3 ± 3.86$^{\dagger\dagger}$ | 29.9 ± 7.98 | 48.8 ± 7.89$^{\dagger\dagger}$ |
| Urine | | | | | | | | | | |
| Diuresis (ml kg-1 BW in 16h) | 2.71 ± 1.07 | 2.57 ± 1.14 | 3.49 ± 1.27 | 6.73 ±1.27$^{\dagger\dagger}$ | 3.01 ± 1.19 | 6.80 ± 0.76$^{\dagger\dagger\dagger}$ | 3.67 ± 1.42 | 8.22 ± 0.72$^{\dagger\dagger}$ | 2.73 ± 1.65 | 7.14 ± 2.30$^{\dagger\dagger}$ |
| Glucose (mg in 16h) | 20.9 ± 10.7 | 46.3 ± 17.4*** | 23.3 ± 5.58 | 27810 ± 2229$^{\dagger\dagger\dagger\dagger}$ | 38.0 ± 17.2 | 26615 ± 1441$^{\dagger\dagger\dagger\dagger}$ | 27.9 ± 7.57 | 3089 ± 2574$^{\dagger\dagger\dagger\dagger}$ | 33.0 ± 12.4 | 28346 ± 3072$^{\dagger\dagger\dagger\dagger}$ |
| Sodium (mg in 16h) | 140 ± 43.3 | 169 ± 30.4 | 160 ± 41.1 | 241 ± 49.5 | 158 ± 57.3 | 312 ± 30.8$^{\dagger\dagger\dagger}$ | 177 ± 63.0 | 268 ± 29.5$^{\dagger}$ | 149 ± 27.7 | 296 ± 60.4$^{\dagger\dagger}$ |
| Creatinine (mg/dl) | 76.8 ± 9.92 | 74.3 ± 12 | 73.2 ± 11 | 82.2 ± 5.58 | 70.2 ± 14.3 | 80.3 ± 3.11 | 71.5 ± 12.0 | 90.4 ± 5.08$^{\dagger}$ | 61.9 ± 7.91 | 85.3 ± 10.9$^{\dagger\dagger}$ |

Values are mean ± SEM. *$p < 0.05$, **$p < 0.01$, ***$p < 0.001$, $^{\#}p < 0.0001$ versus age-matched *Ctns*$^{KO}$ Control and $^{\dagger}p < 0.05$, $^{\dagger\dagger\dagger}p < 0.001$, $^{\dagger\dagger\dagger\dagger}p < 0.0001$ versus *Ctns*$^{KO}$ Dapa. n: number of animals.

**Table S2: Blood parameters in *Ctns* rats after 4 weeks of treatment**

| Plasma | $Ctns^{WT}$ 4 weeks Control (n=6) | $Ctns^{WT}$ 4 weeks Dapa (n=4) | $Ctns^{KO}$ 4 weeks Control (n=6) | $Ctns^{KO}$ 4 weeks Control (n=4) |
|---|---|---|---|---|
| Glucose (mg/dl) | 204 ± 18.4 | 185 ± 6.96 | 205 ± 6.72 | 179 ± 13.6 $^{\dagger}$ |
| Albumin (mg/dl) | 3.66 ± 0.12 | 3.41 ± 0.08$^{\dagger}$ | 3.15 ± 0.12*** | 3.04 ± 0.08 |
| Creatinine (mg/dl) | 0.42 ± 0.10 | 0.50 ± 0.03 | 0.43 ± 0.11 | 0.42 ± 0.03 |
| BUN (mg/dl) | 91 ± 7.77 | 110 ± 13.2 | 85.5 ± 15.1 | 127 ± 12.9 $^{\dagger\dagger\dagger\dagger}$ |
| Total Ketone Bodies (mM) | 5.10 ± 3.15 | 9.16 ± 3.68 | 2.81 ± 2.22 | 8.13 ± 2.04$^{\dagger\dagger}$ |

Values are mean ± SEM. *p < 0.05, ***p < 0.001 versus age-matched $Ctns^{WT}$ Vehicle and $^{\dagger}$p < 0.05, $^{\dagger\dagger\dagger\dagger}$p < 0.0001 versus genotype-matched Dapa. n: number of animals.

**Table S3: Primer sequences**

| **Zebrafish** | | |
|---|---|---|
| Gene | Forward | Reverse |
| *slc5a2* | ATG AGT CGG GTG CTT TCT GG | ATG GCG CAG GGT AAA GAC AA |
| *eef1a1a* | TTC TCC GAG TAT CCT CCT CTG | CTT CTC CAC TCC TTT AAT CAC TCC |

| **Mouse** | | |
|---|---|---|
| Gene | Forward | Reverse |
| *Lrp2* | CAG TGG ATT GGG TAG CAG | GCT TGG GGT CAA CAA CGA TA |
| *Cubn* | TCA TTG GCC TCA GAC ATT CC | CCC AGA CCT TCA CAA AGC TG |
| *Cdc20* | ATG GAG CAG CCT GGA GAC TA | GCT TAC TCG AGC GGA GTG AC |
| *Ccna2* | CTT GGC TGC ACC AAC AGT AA | AGC AAT GAG TGA AGG CAG GT |
| *Ccnb2* | TGA AAC CAG TGC AGA TGG AG | CTG CAG AGC TGA GGG TTC TC |
| *Gapdh* | TGC ACC ACC AAC TGC TTA GC | GGA TGC AGG GAT GAT GTT CT |

| **Rat** | | |
|---|---|---|
| Gene | Forward | Reverse |
| *Slc5a2* | TCT TTG TGC CCC GTG TTA AT | AGT GTA CCC GGC AGA AGA TG |
| *Slc16a1* | GGC AGG GCT ATA AGT CGG TC | GAA TAG AGC AAA GTG CGC GG |
| *Rbp4* | ACT ACG ACA CCT TCG CTC TG | CCT CAG GAA GAG TTG CTC ACC |
| *Baat* | CTG GGG CTA TGA TGA CCT GC | CTG AAG TTA ATG TAA CAC TTG GAC A |
| *Ppard* | AAG GTG TTT CCG GGG TGT TT | TTC GAT CTC TCC ACG CAT CG |
| *Oghd* | TAT GAG CAC AGG TGT CGT CC | CCA GCC AAC CAT TAC CCA GT |
| *Lpl* | GGT GAG CAT CTT GAA CCA AAG C | CGAGCCAGCAGTTGTTTTGA |
| *Acat1* | TCC AAT CGG GTG AGT GCC C | TAA GTC TGT GTT CAC TGC CGG A |
| *Acadl* | AAA CGT CTG GAC TCC GCC TC | TGT CCC TTA TAG ACT GCT TGC TT |
| *Ehhadh* | CCT CAG TTG GCG TTC TTG GT | GAA TGG AGA GAG TTC GTG GG |
| *Acsm3* | AGG ACA GCC TTG AAT ACC TGA GA | AAA TGG GTG TGT GCG TTT CCT |
| *Acot4* | GTC TAT GGT GTT GGA GGG GGC | CAG GAG ACC ACC CCT GTA GA |
| *Acox1* | CCG GCT CCC TTT CTT CTA GG | GCC TAG CACC TGCA CAA AAC TA |
| *Ppargc1a* | GAC CAC AAA CGA TGA CCC TC | TGT TGC GAC TGC GGT TGT |
| *Hrpt* | GCA GAC TTT GCT TTC CTT GG | CAA GGG CAT ATC CAA CAA CA |
